# Quantifying the varying harvest of fermentation products from the human gut microbiota

**DOI:** 10.1101/2024.01.05.573977

**Authors:** Markus Arnoldini, Richa Sharma, Claudia Moresi, Griffin Chure, Julien Chabbey, Emma Slack, Jonas Cremer

## Abstract

Fermentation products released by the gut microbiota provide energy and regulatory functions to the host. Yet, little is known about the magnitude of this metabolic flux and its quantitative dependence on diet and microbiome composition. Here, we establish orthogonal approaches to consistently quantify this flux, integrating data on bacterial metabolism, digestive physiology, and metagenomics. From the nutrients fueling microbiota growth, most carbon ends up in fermentation products and is absorbed by the host. This harvest varies strongly with the amount of complex dietary carbohydrates and is largely independent of bacterial mucin and protein utilization. It covers 2-5% of human energy demand for Western, and up to 10% for non-Western diets. Microbiota composition has little impact on the total harvest but determines the amount of specific fermentation products. This consistent quantification of metabolic fluxes by our analysis framework is crucial to elucidate the gut microbiota’s mechanistic functions in health and disease.

## Introduction

The exchange of fermentation products (mostly acetate, propionate, and butyrate) is a major avenue for the gut microbiota to exert its effects on the host^1,2^. These molecules are excreted by anaerobically growing microbes in the large intestine that feed mostly on complex carbohydrates from plant-based foods, such as dietary fiber and resistant starches, as well as dietary protein that passes the small intestine and mucin^3^. Microbial fermentation products are absorbed by the colonic epithelium and perform a variety of important functions in the host. For example, they serve as an energy source^4,5^ and influence immune cell regulation and recruitment^6–8^, have a role in satiety signaling in rodents^9,10^ and humans^11,12^, and the gut-brain axis more generally^1,13^. By changing the pH in the gut lumen, they also allow gut microbes to shape their local environment with strong effects on microbiota composition^14,15^.

Fermentation product concentrations have been measured in feces and within the human digestive tract^16,17^. However, these concentrations are the result of a highly dynamic interplay between fermentation product release by the microbes and uptake by the host, which varies strongly with changes in the host’s food intake and digestion activity throughout the day^18–20^. Therefore, concentration measurements in gut content are only snapshots and provide little insight into the overall flux of fermentation products that microbes produce and that the human body takes up.

Here, we integrate our own experimental measurements of bacterial fermentation with a quantitative analysis of human digestive physiology to estimate the daily amount of fermentation products the gut microbiota provides to the human host. With this framework, we analyze the variation of the daily fermentation product harvest and its contribution to the host’s energy demand, depending on diet and microbiome composition.

## Results

### Quantifying fermentative metabolism of major gut bacteria

To maintain their redox balance and generate energy in the anoxic environment of the large intestine, gut bacteria utilize different fermentation pathways which mostly produce the acids of acetate, propionate, butyrate, lactate, formate, and succinate (**Figure 1A, Supplementary Text Section 1.1**). To quantify the production of these fermentation products and the consumption of carbohydrates, we grew selected gut bacteria in pure cultures and systematically tracked changes in metabolite concentrations and bacterial biomass over time. Metabolite concentrations change linearly with biomass during steady-state growth, allowing us to determine the per-biomass uptake and excretion rates with high precision (**Figures 1B and S1, STAR Methods, Supplementary Text Section 2.1**).

**Figure 1.**
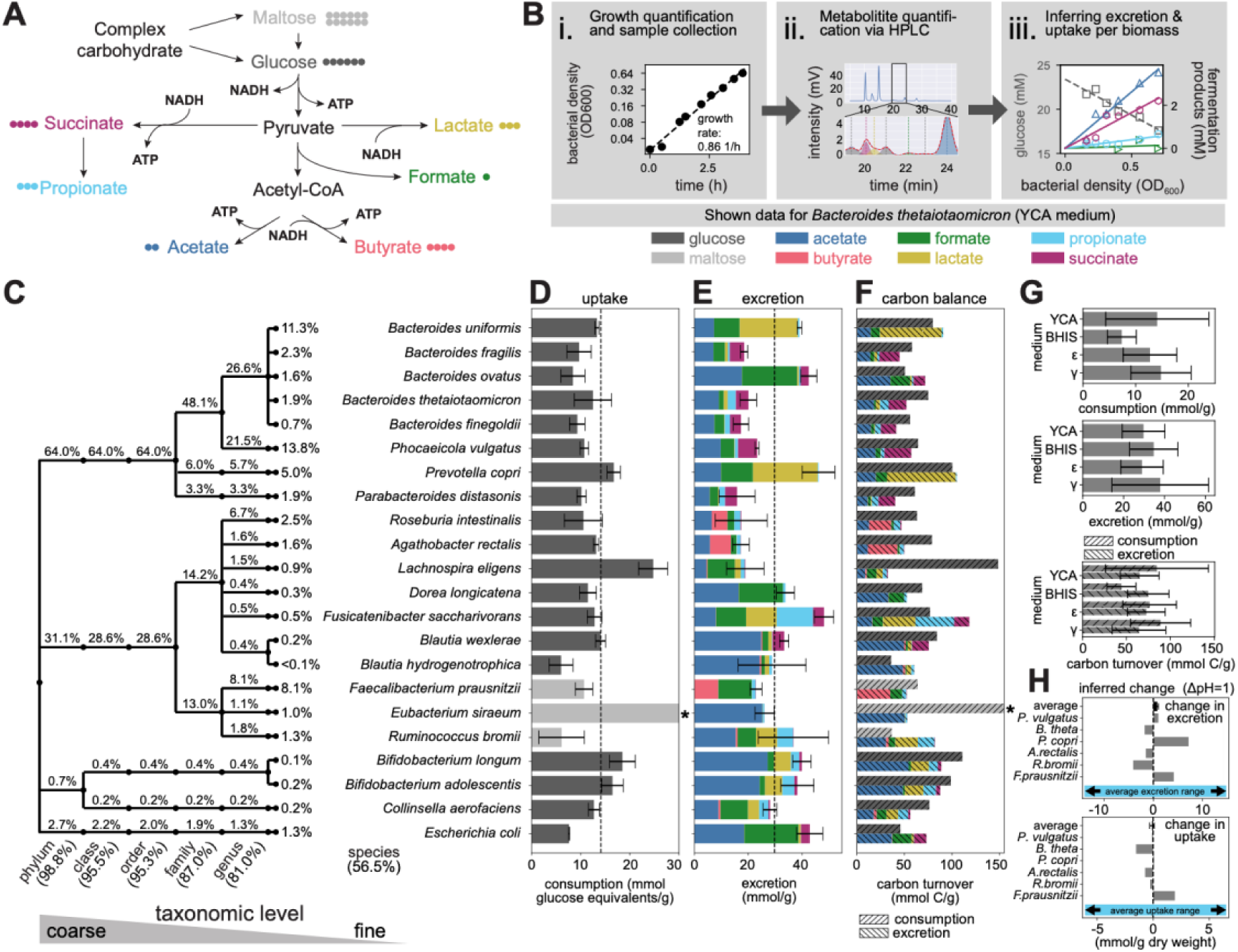
Growth, nutrient uptake, and fermentation product excretion of major gut strains. **(A)** Major fermentation pathways that promote energy synthesis (ATP) and ensure redox balance during anaerobic growth. Colored circles indicate the number of carbon atoms in the corresponding molecules (details on metabolism with full stoichiometry in **Supplementary Text Section 1**). **(B)** Experimental setup to determine nutrient uptake and fermentation product release. Samples taken at multiple time points throughout steady exponential growth (i) were analyzed for metabolite concentration using liquid chromatography (ii). With this data, we calculated linear concentration trends with bacterial density (iii) to quantify per biomass uptake and excretion. Further details in **Figure S1**. **(C)** Taxonomic classification of the 22 gut microbiota species characterized. Numbers indicate in percentage the typical fraction of bacterial biomass these species and their taxonomic groups represent in healthy individuals (samples from^21,22^). **(D,E)** Per biomass glucose/maltose uptake and fermentation product excretion for growth in YCA medium. Error bars denote SD of three biological replicates for sugar update and total fermentation product secretion, respectively. Dashed lines indicate averages of all measured species. **(F)** Balance of carbon content in taken-up carbohydrates and excreted fermentation products. Mismatched carbon balances for *E.* coli, *R. bromii*, and *F. saccharivorans* likely reflect the utilization of other media components as carbon sources; *for B. hydrogenotrophica* the imbalance is likely due to acetogenesis. For *E. siraeum*, carbohydrate uptake is higher than the range shown (27.3 mmol maltose/g, stars in D,E). **(G)** Species averaged uptake and excretion rates in different media (YCA, BHIS, 𝜖, 𝛾). Error bars denote SD of species-to-species variation. **(H)** Dependence of fermentation product excretion and carbohydrate uptake on pH. Changes for shifts in pH by 1 unit are based on linear regressions of measured values (**Figure S3DE**). Range of the horizontal axes were chosen to reflect the average uptake and excretion ranges determined in (**D, E**)

We selected 22 bacterial species (**Figure 1C** and listed in the **Key Resources Table**) based on their relative abundances in a typical healthy human gut microbiota. While hundreds of species are usually detectable in a gut microbiome, their abundances are highly uneven, with the selected species commonly accounting for close to 60% of total bacterial biomass in the healthy human gut (**Figure S2**, based on 219 samples from healthy adults^21,22^). When using these species as representatives of their respective genera, coverage increases to around 84% of biomass (**Figures 1C, S2**). For most tested species, including those most prevalent in the gut, uptake of glucose or maltose varied in a relatively narrow range (**Figure 1D**). In contrast, the types of secreted fermentation products varied substantially between different bacterial species, reflecting their different metabolic pathways (**Figure 1E**). For example, *Bacteroides* strains excrete succinate while many abundant Lachnospiraceae excrete butyrate (**Supplementary Text Section 1.1**). We found little variation in the fate of carbohydrate-derived carbon, most of which ends up in fermentation products (**Figure 1F**), illustrating the high demand for carbohydrates required for fermentative growth.

In the gut, nutrient availability and pH conditions can vary strongly^18,23,24^. To test how robust our results are beyond growth in the rich YCA medium discussed before, we first repeated these experiments in media of different complexities (BHIS, 𝜖, and 𝛾 media; see **STAR Methods** for details). Per-biomass carbohydrate uptake and fermentation product excretion are similar across media (except for carbohydrate uptake in BHIS, which we likely underestimate as it contains carbohydrates not captured by our measurement assay), see **Figures 1G** and **S3A-C**. Second, we have probed uptake and excretion characteristics of selected highly abundant strains for a range of pH conditions relevant in the large intestine (5.5-7.5)^23^. While growth rates can vary, we see only minor changes in uptake and excretion per-biomass (**Figures 1H** and **S3D-H**). This finding indicates that bacteria face the same fundamental metabolic constraints across conditions, and the experimentally determined values therefore provide robust estimates. This robustness can be further rationalized by the ATP requirement of growing microbes, which is dominated by biomass synthesis and does not change much with different conditions (**Supplementary Text Section 1.2**). In the following analysis, we use the uptake and excretion values measured in YCA medium.

### Point estimates of the daily fermentation product harvest

Integrating our experimental dataset and a quantitative analysis of diet and digestion, we next estimated the daily amount of fermentation products released by the gut microbiota in two different ways. To do so, we use measurements performed in 1970’s Great Britain, where dietary and digestion data has been reported in unparalleled quality. First, we estimated the fermentation product release based on fecal weight (**Figure 2A and Supplementary Text Section 3.1**): For the British reference scenario, humans excrete around 30g of fecal dry weight every day^25^ (**Figure 2A(i), Figure S4A-D**) of which around 16g is bacterial biomass^26^ (**Figure 2A(ii)**). To maintain a stable microbiota in the gut, this loss of bacterial biomass must be compensated by bacterial growth. Using the measured amount of fermentation products that bacteria release to support growth and weighting them by the average genus-level abundance in the fecal microbiome samples from healthy adults (**Supplementary Text Section 2.2**), we get a daily production of fermentation products of approximately 16g/day x 29mmol/g ≈ 470mmol/day (**Figure 2A(iii)**). Second, we estimated the fermentation product release based on the amount of carbohydrates in the diet (**Figure 2B and Supplementary Text Section 3.2**). From the amount of fiber, sugar, and other carbohydrates in the diet and their digestibility, we estimated the amount of carbohydrates reaching the large intestine, of which a substantial fraction can be utilized by the gut microbiota (the microbiota available carbohydrates, **Supplementary Text Section 4**). For the British reference scenario, the microbiota available carbohydrates amount to 36g/day^27^ (**Figures 2B(i) and 2C**, corresponding to 198mmol equivalents of glucose), with exact values depending on the starch and fiber content in the diet (**Figure S4E-J**, **Supplementary Text Section 4**). Using the measured amount of carbohydrate needed to produce bacterial biomass (on average 13 mmol glucose equivalents/g biomass,) approximately 198mmol / 13mmol/g ≈ 16g bacterial biomass is produced per day (**Figure 2B(ii)**), leading to a daily fermentation product release of 16g * 29mmol/g ≈ 450 mmol (**Figure 2B(iii)**), in good agreement with the estimation via fecal weight. When estimating fermentation product release based on the experimentally measured uptake and excretion for other media, we arrive at similar results (**Figure S5A**), highlighting again the consistency of the estimations independent of growth conditions.

**Figure 2.**
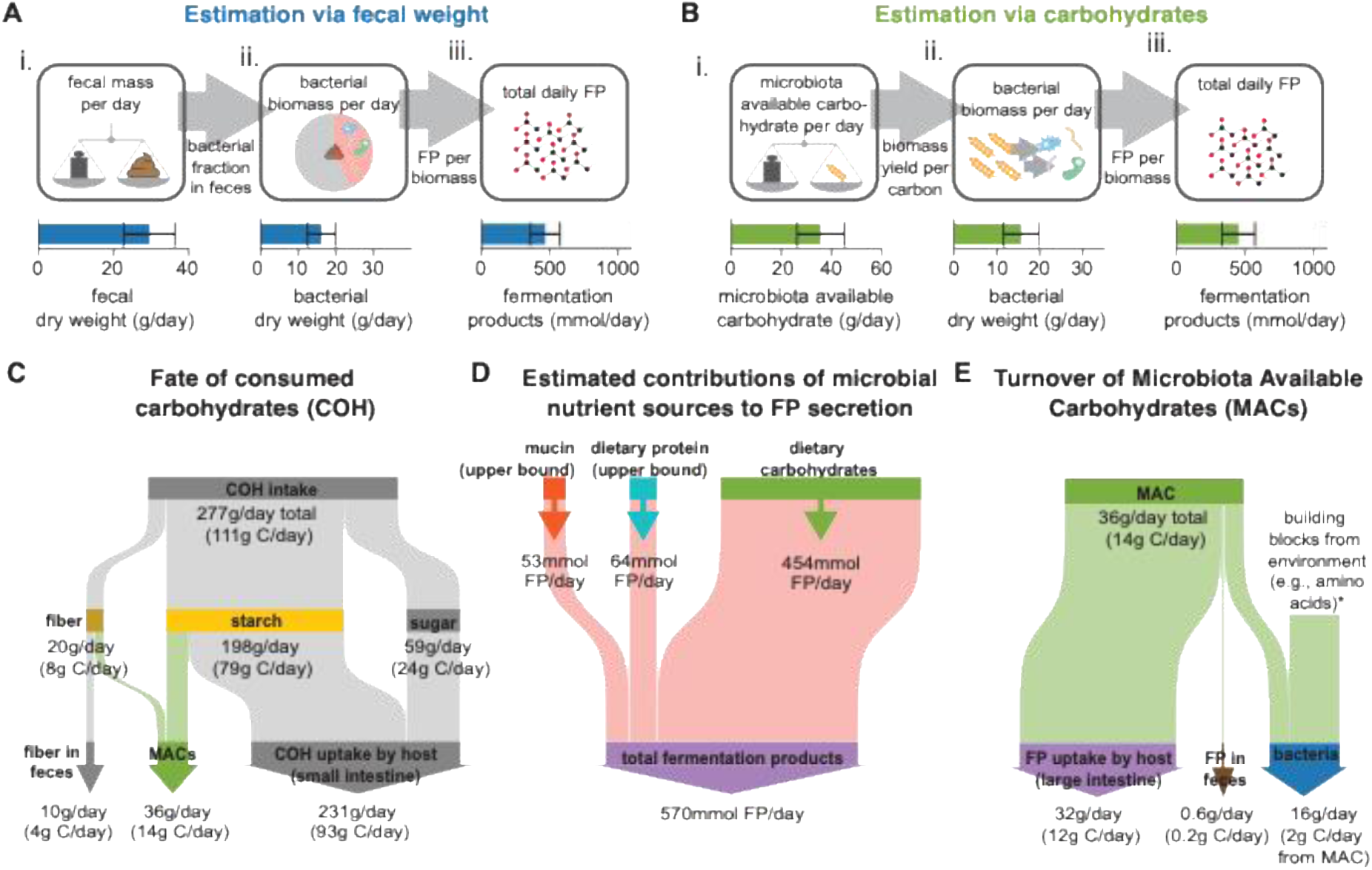
Daily fermentation product harvest for British reference scenario. **(A)** Fecal-weight-based point estimate of the total daily fermentation product release. Starting point is the measurement of daily fecal weight^25^ (i). Given the fraction of bacteria in fecal weight^26^, bacterial biomass in feces follows (ii) which, given the measured fermentation product excretion, sets the total daily fermentation product release (iii). **(B)** Carbohydrate-based point estimate of the total daily fermentation product release. Starting point is the estimation of microbiota available carbohydrates (i). With experimentally determined uptake and excretion rates (Figure 1), the bacterial biomass that gut bacteria synthesize (ii) and the resulting fermentation product harvest when bacteria grow on these carbohydrates (iii) follows. **(C)** Flow diagram showing the daily turnover of consumed carbohydrates along the upper digestive tract with only a fraction being available for the microbiota (details of this mapping described in **Supplementary Text Section 4**). **(D)** Contribution of bacterial mucin and protein degradation to fermentation product release. Compared to dietary carbohydrates, contributions are much smaller, even when systematically overestimating them (**Supplementary Text Section 5**). **(E)** Flow diagram showing resulting carbohydrate and carbon flow along the large intestine, with most of the carbon ending up in fermentation products that are absorbed by the host. Calculations are based on measured uptake and excretion rates in YCA medium (Figure 1D-E) weighted by genus abundance (**Supplementary Text Section 2.2**). g C/day of total carbohydrates, fiber, starch, sugar, and microbiota-available carbohydrates were calculated by assuming a fraction of carbon per total weight equal to glucose (0.4); the fraction of carbon in bacterial biomass was taken to be 0.39, based on data in^71^. Corresponding plots based on experiments in other media are shown in **Figure S5**. Error bars in A and C denote SD based on the variation in daily fecal weight and microbiota-available carbohydrates, respectively. Calculations described in **Supplementary Text Section 3**.

In addition to dietary carbohydrates, a fraction of dietary protein and host-derived mucin serve as nutrient sources for bacteria in the colon^28,29^. We have quantified the contribution of these nutrient sources to the daily release of fermentation products by the gut microbiota, accounting for dietary protein consumption and digestion, mucin turnover, bacterial protein degradation, and amino acid metabolism (**Supplementary Text Section 5**). Even though we consistently used upper bound estimations in our analysis (e.g., all amino acids are fermented and not used for the synthesis of new bacterial proteins), utilization of peptides and mucin contribute a maximum of 20% of the total fermentation product release (**Figure 2D**, **Supplementary Text Section 5**), with the actual number likely being much smaller. Thus, diet-derived carbohydrates account for the vast majority of the microbiota’s nutrient supply. This is further supported by the consistency between our two estimates of total fermentation product release based on fecal weight or dietary carbohydrates.

Given the high carbohydrate demand of anaerobic catabolism, most of the carbon from microbiota-available carbohydrates will end up in fermentation products (>90%, **Figure 2E**). Less than 2% of these exit the host via feces^30^, (**Figures 2E and S5B,C**, **Supplementary Text Section 6**). As the effect of cross-feeding on total fermentation product levels is likely limited (**Supplementary Text Section 6**), the daily amount of fermentation products released by bacteria represents a good upper bound for the daily amount absorbed by the gut epithelium and thus, the daily fermentation product harvest by the human host.

### Variation in total fermentation product harvest with changing microbiota composition

For the point estimates of the daily fermentation product harvest presented above, we assumed a constant microbiome composition typically found in healthy humans. However, gut microbiota composition varies. For example, the relative abundance of primary fermenters from the families Bacteroidaceae and Lachnospiraceae varies considerably between the 219 mentioned microbiome samples from healthy adults (**Figure 3A**, upper panel)^22^. To illustrate the effect of these changes in microbiota composition on fermentation product harvest, we weighted measured consumption and production rates of different species with their relative abundances in the microbiome samples, using them as metabolic representatives of their respective genera (**Figure S6**, **Supplementary Text Section 2.2**). We kept all other parameters constant, including the daily supply of microbiota available carbohydrates, for which we here use the values for the British reference scenario. The relative amount of different fermentation products released varies substantially with microbiome composition (**Figures 3A** lower panel, and **B**). However, both the daily production of bacterial biomass (**Figure 3C**) and the total daily release of fermentation products (**Figure 3D**) show less change. These differences in variation are quantified by the coefficients of variation (**Figure 3E**), with lactate and butyrate showing the highest values. Changes in fermentation product release with host age, health, and lifestyle, obtained from a collection of published metagenomic data (93 studies with about 18000 samples in total covering various health conditions, ages, and lifestyles)^31^ can be explored in an interactive figure (**Data S2** or https://cremerlab.github.io/fermentation_products/study-explorer) accompanying this paper. In summary, this analysis shows that, given a specific diet, differences in microbiota composition lead to differences in the types of fermentation products released, while the total daily release of fermentation products remains within a narrow range.

**Figure 3.**
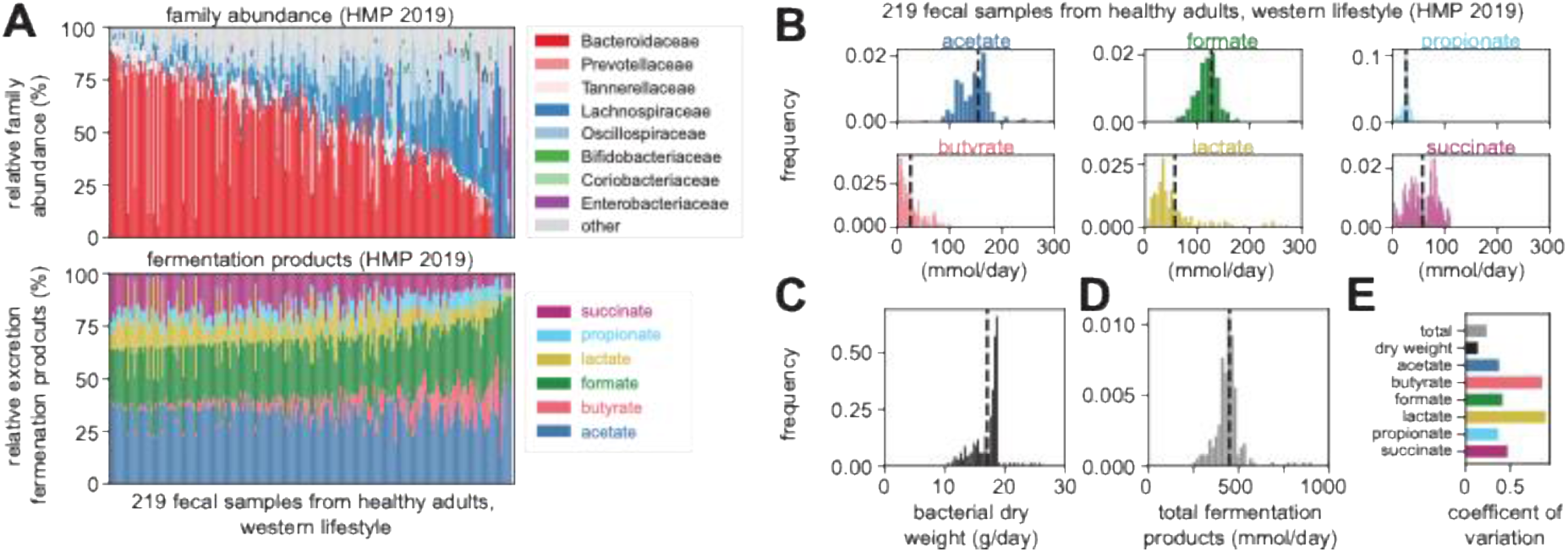
The effect of microbiota composition on daily fermentation product harvest. **(A)** Variation of gut microbiota composition on the family level (top) and corresponding estimated molar fraction of different fermentation products (bottom) across fecal samples from healthy individuals^21,22^. **(B, C, D)** Distribution of the daily release of single fermentation products (**B**), total bacterial biomass (**C**), and total amount of fermentation products (**D**) for the same set of samples. **(E)** Coefficients of variation quantify these variations, with the variation in total fermentation product release being much smaller than variations in the production of specific fermentation products. Estimations for a fixed amount of microbiota available carbohydrates following the 1970’s British reference scenario (Figure 2**B, C**). Calculations described in **Supplementary Text Section 2 and 3**. The variation with carbohydrate consumption and for further metagenomics datasets can be explored in an interactive figure (**Data S2**).

### Variation in total fermentation product harvest with changing diet

To quantify the variation in daily fermentation product harvest with diet, we next analyzed data on diet composition and digestion from three different cohorts, covering much of the global variation in human diets.

First, we estimated the variation in the US population using nutrition data from the 2017/2018 National Health and Nutrition Examination Survey (NHANES) ^32^ (**Figure S7A-F**). We inferred inter-individual variation in daily intake of microbiota-available carbohydrates based on nutrition data in this cohort (**Figure S8A-C**) using the same mapping as above (**Figure 2C, Supplementary Text Section 4**). We then estimated the fermentation product harvest via carbohydrates using this variation while keeping microbiome composition constant (using abundance numbers for a typical healthy human gut microbiome, analogous to **Figure 2BC**). The fermentation product harvest is broadly distributed around a median of 286mmol/day (**Figure 4A**), with most individuals staying substantially below the estimate for the British reference scenario (purple dashed line), reflecting how modern Western diets lead to a lower fermentation product harvest from the gut microbiota^33–35^. The impact of protein fermentation based on protein consumption is again small, **Figure S8D**.

**Figure 4.**
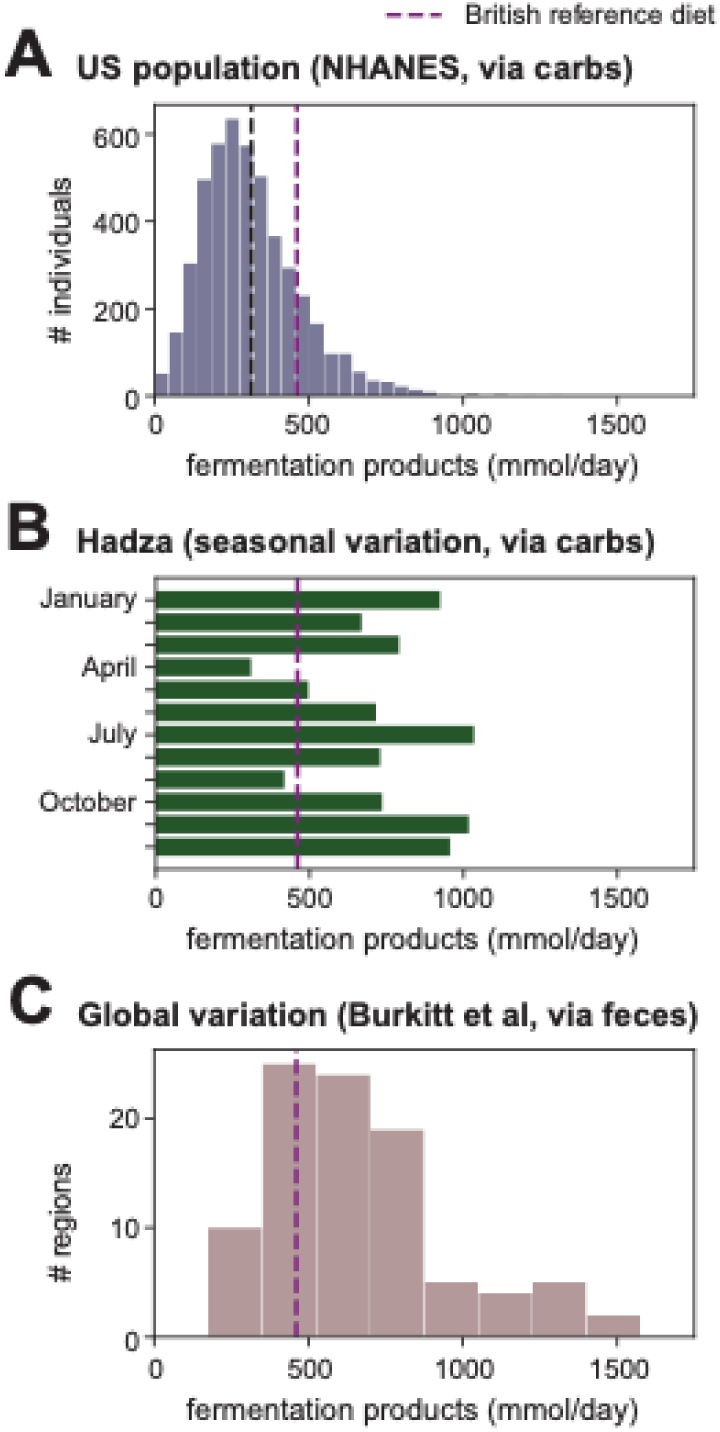
Variation of the daily fermentation product harvest with diet and digestion. **(A)** Inter-individual variation in total daily fermentation product release for the US population. Carbohydrate-based estimate using reported variation in diet for the NHANES 2017/2018 cohort^32^. **(B)** Month-to-month variation of total daily fermentation product release for the Hadza. Carbohydrate-based estimate using reported seasonal variation in diet^36^. **(C)** Variation in fermentation product release based on fecal-based estimate and global variation in fecal weight reported in Burkitt et al^38^. These numbers represent an upper bound estimation (**Figure S9** for detailed discussion). Shown estimates are based on per-biomass uptake and excretion rates for the 22 strains characterized in Figure 1, weighted by their genus-level abundance in a typical healthy human microbiome.

Second, we analyzed nutrition data available for the Hadza, an indigenous hunter-gatherer group in Tanzania. Their diet, including the amount of carbohydrates consumed, differs substantially from a modern Western diet, has strong seasonal variation, and often includes fiber-rich tubers^36,37^ (**Figure S7G, H**). Accordingly, the estimated amount of microbiota-available carbohydrates (**Figure S8E-G**) and the resulting daily fermentation product harvest show strong seasonal variation (**Figure 4B, Supplementary Text Section 4**; protein fermentation has again a minor effect on total fermentation product harvest, **Figure S8H**). We find that, for the Hadza, harvests are much higher than those typically observed for people consuming Western diets and can reach up to approximately 1000mmol/day with strong seasonal variations.

Third, we estimated the global variation in fermentation product harvest based on fecal weight data representing different geographical regions and lifestyles around 1970^38^, before Western diets became more prevalent around the world. Daily excretion of fecal wet weight varies strongly, with maximum values about five times higher than in the British cohort (**Figure S9A**). The resulting variation in daily fermentation product harvest is shown in **Figure 4C**, and reaches a maximum value of 1500mmol/day. For very high fecal mass, this is likely and overestimation: while we corrected for changes in water content with fecal mass^17,26,39–43^, we keep the bacterial fraction per fecal dry weight constant due to a lack of available data (**Figure S9**). As differences in fecal weight are largely due to differences in diet^39,43,44^, this broad variation again emphasizes the importance of diet for the fermentation product harvest. In summary, we find that diet, and not microbiota composition, is the dominant factor in setting this number. Particularly, we show how diets low in microbiota-available carbohydrates, as consumed in the US and increasingly around the globe, exhibit much lower fermentation product yields than fiber-rich diets, with potentially detrimental health outcomes^33–35^. Variation of fermentation product harvest with diet, as well as its interaction with changing microbiome composition, can furth be explored in the **Data S2**.

### Microbiota-derived fermentation products as energy source for the host

We next compared the human fermentation product harvest with that of the mouse, the most common experimental model in gut microbiota research. Hoces and co-workers have recently compared the energy extraction from food in germ-free and conventionally colonized mice ^19^. Energy extraction is lower by 8.1kJ/day in germ-free mice (**Figure 5A,B**), likely due to a lack of microbiota-available-carbohydrate-derived fermentation products that serve as energy sources for the host (**Supplementary Text Section 7.1**). When estimating the harvest of fermentation products (via fecal weight, analogous to **Figure 2A**) and their energy content *(i.e.,* combustion enthalpy, see **Supplementary Text Section 8, Table S1**) for these mice, we find that fermentation products can explain this difference in energy extraction (**Figure 5C**, **Supplementary Text Section 7.2**), indicating that most fermentation products are utilized as an energy source by the host. In the same way, we calculated the energy content of the fermentation product harvest in humans. Depending on diet and digestion, these values vary between 0.1MJ/day and 1.2 MJ/day. To put these numbers in perspective and allow a comparison with values in mice, we normalized them by the daily energy expenditure of the respective host (approximately 10MJ/day for humans^45^ and 38kJ/day for mice^19^). For the British reference scenario, the gut microbiota contributes 4.4% of the host’s daily energy expenditure (**Figure 6**, purple dashed line). We find broad variation around this number, with values varying from 1.7% (5th percentile of the NHANES cohort, **Figure 6**, blue) to 12.1% in the extreme cases of non-Western diets (**Figure 6**, red; 95th percentile from Burkitt data). In contrast, for mice, we arrive at 21% or more of the total energy demand of 38kJ/day^19^ (**Figure 6**, grey, and **Supplementary Text Section 7**). Notably, these mice were fed autoclaved laboratory chow, and mice consuming non-autoclaved food or a more natural diet are likely to consume more microbiota-available carbohydrates, resulting in an even higher microbiota-derived energy supply^46^. Therefore, our analysis quantifies a substantial difference in microbiota-supplied energy between humans and mice.

**Figure 5.**
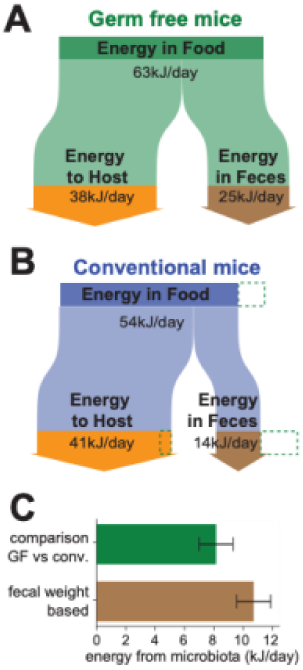
Daily metabolic energy contribution of the gut microbiota in mice. **(A, B)** Flow charts illustrate the flow of energy consumed as food in germ-free (**A**) and conventionally colonized mice (**B**). Around 60% or 75% of the consumed energy is extracted in germ-free and conventionally colonized mice, respectively. The rest is excreted in feces. Accordingly, the energy consumed in food and energy in feces is lower in conventional mice, while the energy retained in the host is slightly higher. Green dashed lines in B indicate these differences. **(C)** Daily host energy provided by the gut microbiota when comparing the energy retention in germ-free and conventional mice (upper bar) or when estimating fermentation product release using fecal mass in mice (lower bar, analogous to Figure 2A). Error bars denote SD based on the variation of consumed food and fecal weight of different mice (upper estimation). Calculations and variation estimates are detailed in **Supplementary Text Section 7**. Based on data reported in^19^.

**Figure 6.**
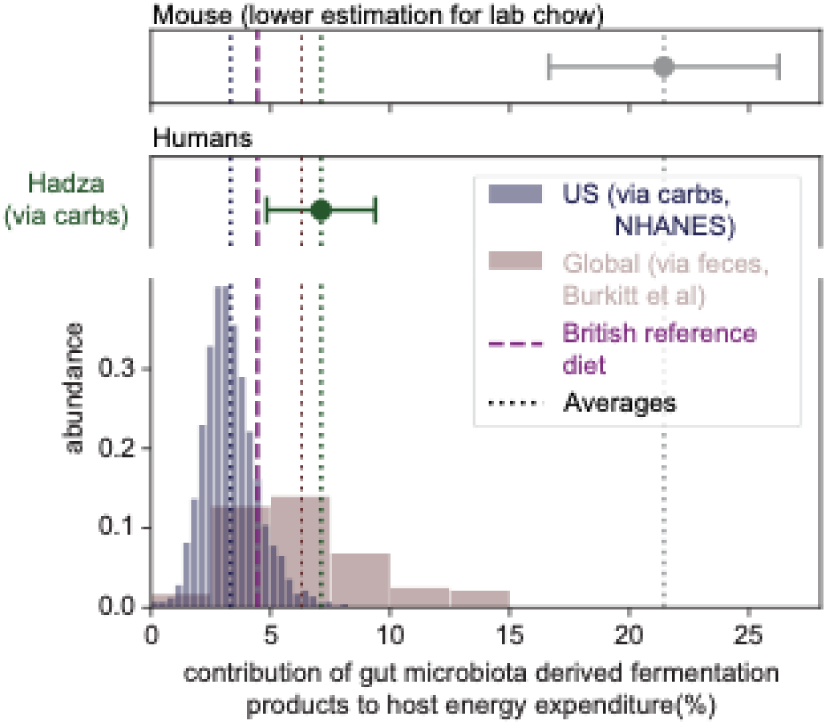
Microbiota contribution to energy supply. Variation of fermentation product harvest leads to strong variation in the fraction of daily energy demand supplied by the gut microbiota. Non-Western diets show much higher values (green, red), with an upper bound estimation of >10%, when compared to a cohort of US residents (blue). Laboratory mice feeding on autoclaved lab chow (grey) show a substantially higher percentage of daily energy covered by the microbiota. Values in mice are likely even higher when they consume non-autoclaved food reflecting their natural diet. Error bars indicate variation (SD) based on monthly variation in food intake (Hadza data, green, based on data in^36^), and on error propagation using data originally published in^19^ (mouse data, gray, see **Supplementary Text Section 7.2**), respectively.

## Discussion

Recent studies have emphasized the importance of quantitative physiological considerations in microbiome research^14,47,48^. By integrating data on nutrient intake and fecal mass with experimental measurements of bacterial growth and fermentation rates, we quantified the daily fermentation product harvest in the human intestine.

Notably, complementary analyses led to similar outcomes, confirming the validity of our integrative framework. First, the fermentation product harvest is highly comparable when deriving it from consumed carbohydrates or released feces, both for the point estimate of the British reference scenario and when analyzing its variation (**Figures 2A,B and 4**). Second, a revised theoretical consideration based on the ATP turnover required to generate bacterial biomass^49^ predicts a similar total fermentation product harvest for the British reference scenario (**Supplementary Text Section 3.3**). Third, estimating the fermentation product harvests via fecal weight matches an analysis of energy extraction by the microbiota in mice (**Figure 5C**).

When utilizing this framework to analyze fermentation product harvest, several findings stand out: the bulk of carbon in microbiota available carbohydrates, >90%, ends up in fermentation products, most of which are taken up by the host (**Figure 2E**). Quantitatively, fermentation products, therefore, dominate the metabolite exchange between the microbiota and the host, affecting host signaling, behavior, and energy homeostasis. Differences in the total fermentation product harvest are primarily determined by variation in diet. Low fermentation product harvests could result from the consumption of highly processed foods low in complex carbohydrates, or of food that contains more resistant carbohydrates which leave the body undigested. In contrast, microbiota composition has less impact on the total daily fermentation product harvest. However, microbiota composition strongly impacts the harvest of specific fermentation products, with butyrate and lactate showing the strongest variation with composition (**Figure 3E**). Given the different ways specific fermentation products affect host processes^50^, this finding may underlie many of the commonly reported links between health status and microbiota composition, even without explicitly considering the dominant effect of diet on the total fermentation product harvest. However, as dietary changes also cause changes in microbiota composition^51^, diet is likely the key determinant for both, the abundance of specific fermentation products and the total fermentation product harvest. As such, our results highlight that the axis between diet, microbiota composition, and fermentation product harvest should routinely be considered when investigating host-microbiota interactions, their role in host health, and the etiology and progression of different microbiota-associated diseases. Critically, because the vast majority of fermentation products are taken up locally in the large intestine, fermentation products measured in feces represent only a small fraction of the total production. Total fermentation product release can be better estimated with the integrative framework presented here based on diet, microbiome composition, and the constraints of bacterial fermentation.

Our analysis also reveals an important systemic difference between humans and mice. Specifically, the fermentation product harvest per body weight is much higher in mice than in humans (approximately 400 mmol/kg/day for the mouse vs. 7 mmol/kg/day for humans, **Supplementary Text Section 7.3**). This difference is also reflected in the contribution of microbiota-derived fermentation products to the daily energy demand, which is much higher in mice compared to humans (1.7-12.1% in humans vs more than 21 % in laboratory mice, **Figure 6**). Together with previously reported differences in microbiome composition and anatomy of the digestive tract^52–56^, this difference in fermentation product harvest needs to be accounted for when utilizing mouse studies to explore possible systemic effects of the microbiota on the human host. For example, while specific microbiota perturbations have strong physiological and behavioral effects in mice, comparable perturbations in humans might have a smaller impact^57^, simply because mice rely more on fermentation product-derived energy than humans. On the other hand, local characteristics like the absorption of fermentation products per epithelial surface area might be more comparable (170 and 220 mmol/m^2^/day in mice and humans, respectively; **Supplementary Text Section 7.3**), suggesting that local interactions between the human microbiota and the intestinal mucosa can be emulated more realistically in mouse studies.

More generally, this study emphasizes the strong potential of quantitative analyses that incorporate microbial metabolism, diet, host physiology, and microbiota turnover to put numbers on the metabolic exchange between microbiota and host. By providing information on specific doses, this goes well beyond the identification of microbiota-derived molecules and their possible interactions with the host. Future studies using such quantitative insights on metabolite fluxes in clinical settings or to inform genome-scale metabolic models^58,59^ can be instrumental in establishing mechanistic links between microbiome composition, the exchange of specific fermentation products, and the onset and progression of different diseases.

In addition to the quantification of major fermentation products and their harvest in this study, many other interaction paths would benefit from such an integrative analysis. This includes the nitrogen cycling between the host and the gut microbiota^60^, the role of cross-feeding in the exchange of gases such as H_2_ ^61^ and CO, and the microbial digestion of proteins which exposes the host to branched-chained fatty acids and toxic waste products like hydrogen sulfide. In the context of the dynamic, flowing environment of the intestine^62^, developing such frameworks is critical to move the field away from assumptions based on point measurements of metabolite concentrations towards an integrated model of host-microbiome functions driving health and disease.

## Limitations of the study

To better understand microbiome-host interactions, we focus on the fermentation product harvest, quantitatively the largest flux of microbial metabolites reaching the host. Our approach has three major limitations which should be addressed in future studies. First, our analysis framework uses a static microbiome composition as input. As such, it cannot predict changes in microbiome composition and, thus, changes in fermentation product turnover over time. Implementing this would be a highly complex problem due to the dynamic eco-physiological interactions between hundreds of microbial species and the host, which strongly depend on pH^14,15,63^, transit times^64,65^, the carbohydrate preferences of different microbes^51^, and other parameters of the system. Second, we did not explicitly consider metabolite cross-feeding in this study. Most cross-feeding processes would not change the total abundance of fermentation products, but convert them into other fermentation products. Therefore, cross-feeding can change the abundance of specific fermentation products (**Supplementary Text Section 6**). While our experimental data implicitly accounts for cross-feeding of acetate, further work on the quantitative role of cross-feeding on lactate is needed to better understand the molecular composition of the fermentation product harvest, especially in microbiomes where the relative abundance of microbiota members performing this function is high^66–68^. Third, we focus on the central metabolites of fermentation, accounting for most of the molecules exchanged between the microbiota and host. However, other, less abundant chemicals, such as H_2_S^69^ or TMA^70^, could have significant and potentially detrimental consequences on the host. For all three points, further quantitative studies are needed. Our present analysis is a starting point for such efforts since fermentation is the major source of energy which other microbial processes in the gut depend on.

## Resource Availability

### Lead Contact

Further information and requests for resources including data and coding scripts, should be directed to and will be fulfilled by the lead contact, Jonas Cremer.

### Materials Availability

No new unique reagents or strains were generated in this study. Complete information of all materials used in the study can be found in the Key Resources Table.

### Data and Code Availability

Customized Python scripts, raw experimental data (including chromatography data and growth curves), as well as processed data, are available on the paper’s GitHub repository (https://doi.org/10.5281/zenodo.10445504), accessible via https://github.com/cremerlab/fermentation_products.

## Acknowledgements

We thank Daniel Hoces, Verena Lentsch, Anna Sintsova, Alice de Wouters, Alfred Spormann, and all members of the Cremer, Spormann, and Slack groups for suggestions and discussions. We are grateful to Kevin Foster, Wolf-Dietrich Hardt, Ron Milo, and Uwe Sauer for critical reading and comments that helped improve the manuscript. MA and ES were supported as a part of NCCR Microbiomes, a National Centre of Competence in Research, funded by the Swiss National Science Foundation (grant number 180575), and by project grants from the Swiss National Science Foundation (grant numbers 40B2-0_180953, 310030_185128). CM was supported by an Innosuisse grant (grant number 59256.1 IP-LS). GC was supported by the NSF Postdoctoral Research Fellowships in Biology Program (grant number 2010807). ES was also supported by the Botnar Research Centre for Child Health as part of the Multi-Investigator Project: Microbiota Engineering for Child Health. JC acknowledges support by the National Institute of General Medical Sciences, National Institutes of Health (grant number 1R01GM149611), a Stanford Bio-X Seeding Grant (grant number 10-32), a Center for Pediatric IBD and Celiac Disease Seed Grant (309906), and a Terman fellowship.

## Author contributions

Conceptualization, MA, JCr; methodology, MA, GC, JCr; investigation, MA, RS, CM, JCh, JCr; formal analysis, MA, JCr; software, GC, JCr; visualization, MA, JCr, GC; data curation, JCr; funding acquisition, MA, ES, JCr; resources, MA, ES, JCr; writing, original draft, MA, JCr; writing, review & editing, all authors

## Supplemental Figures

**Figure S1.**
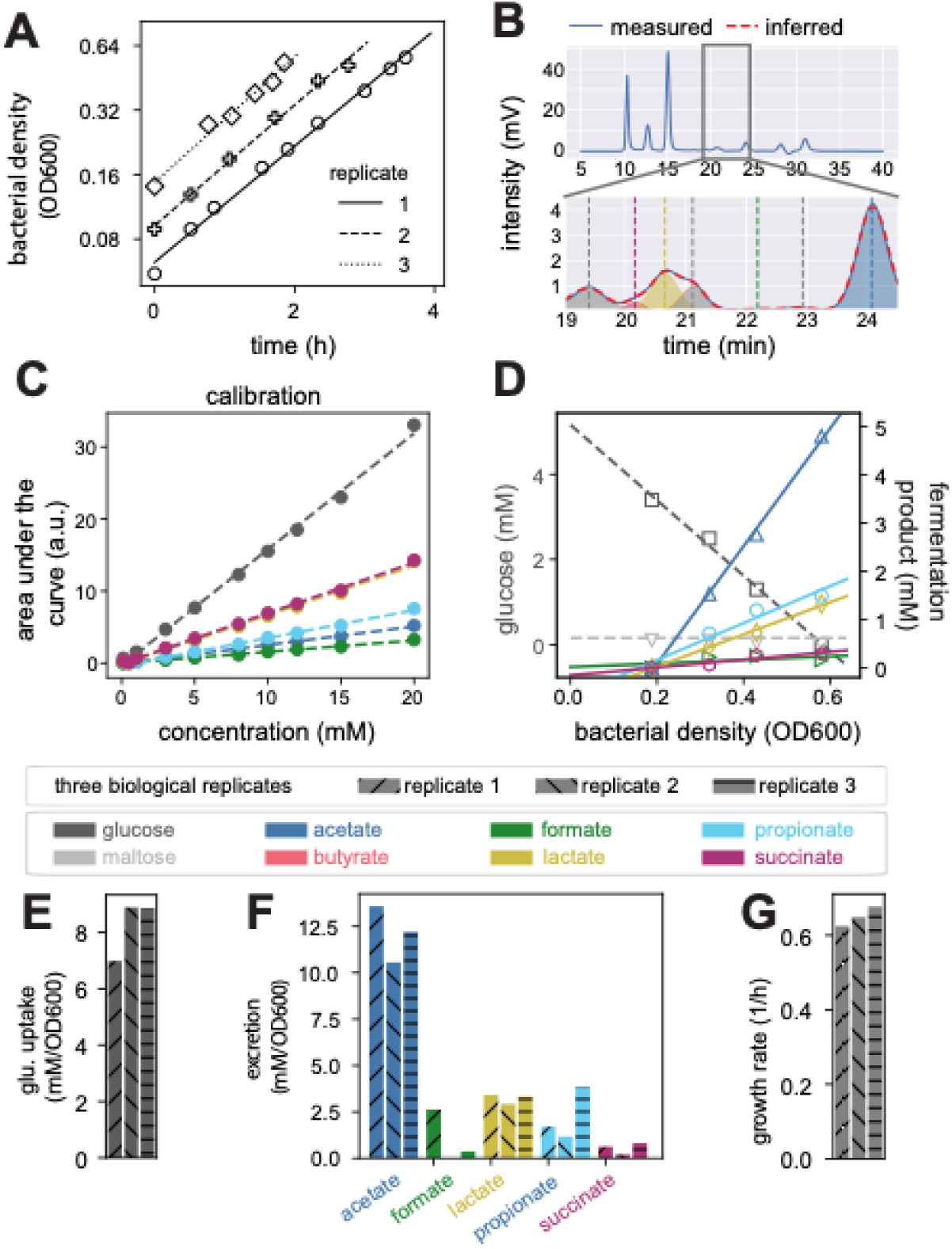
Experimental workflow to determine the per-biomass excretion and consumption rates of metabolites, Related to Figure 1. (**A**) Optical densities (OD600) were measured over time and samples were taken at different OD600 values during exponential growth (symbols). Culturing in two steps included a pre-culture phase before sample taking to ensure steady growth. Different lines show three independent biological replicates. (**B**) These samples were then analyzed for metabolites using HPLC. Peaks for specific metabolites were identified, computationally isolated, and the areas under the curves they cover were quantified using a Python-based data analysis pipeline we have developed^14,72^. (**C**) Obtained peak areas were subsequently compared to the shown component-specific standard curves to determine their concentrations. (**D**) Linear fits describing the change of these concentrations with optical density, as expected for steady growth, were then used to calculate the per biomass production and consumption of different metabolites. (**E, F**) Per-biomass excretion of fermentation products, {𝑒_𝑖_}, and uptake of glucose, 𝑢, for three biological replicates. (**G**) Exponential growth rates 𝜇 of three biological replicates, obtained by a linear regression on the log-transformed OD data shown in (**A**). From these numbers, excretion and uptake per time can also be calculated. Exemplary data shown here for *B. theta* growing in YCA medium. Similar plots for other species and growth conditions are available via the GitHub repository (folder hplc_measurements).

**Figure S2.**
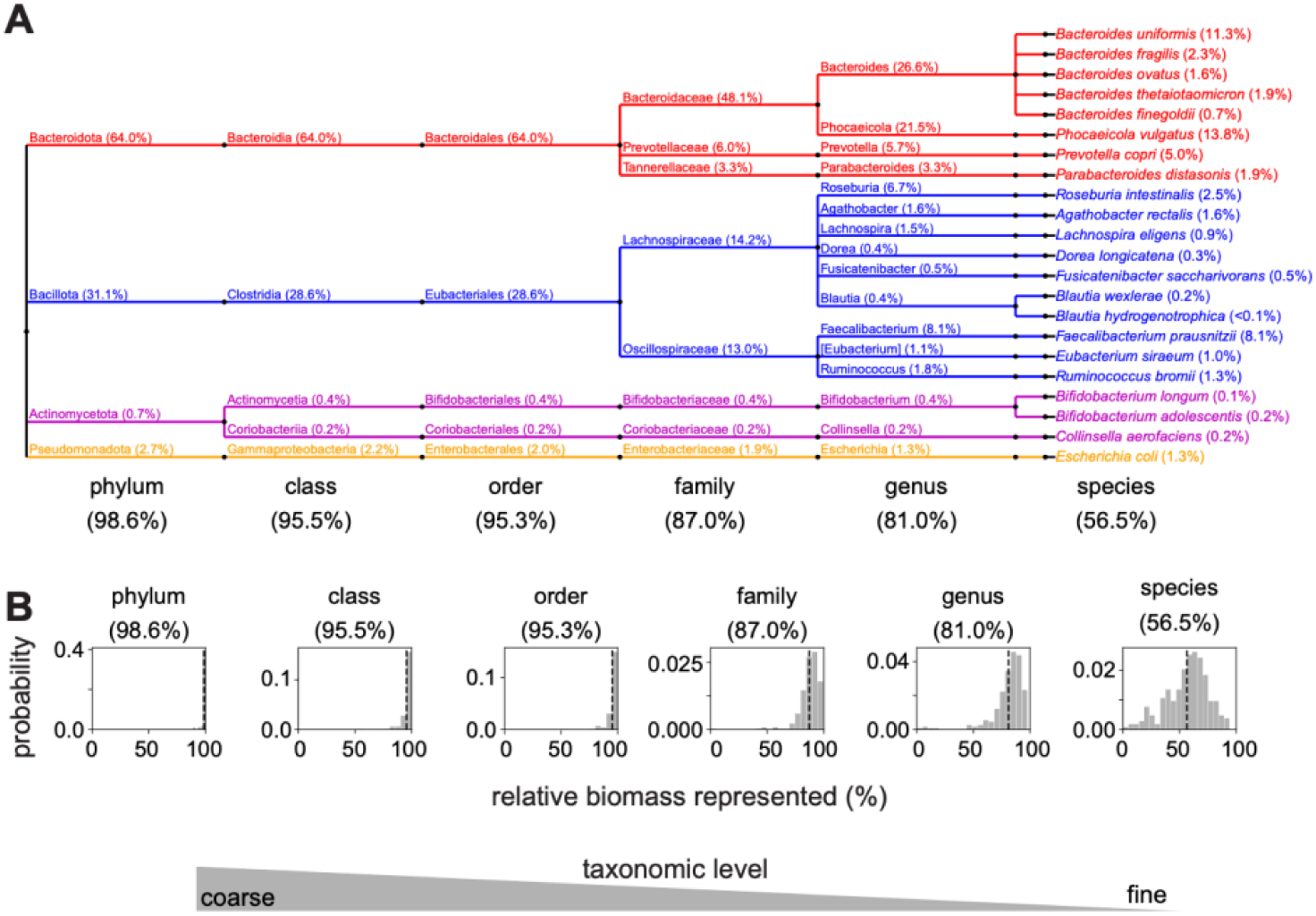
Taxonomic relation and biomass coverage of experimentally characterized strains, Related to Figure 1. (**A**) Phylogenetic tree of 22 highly abundant human gut microbiota strains we characterized. Branch labels indicate the names of different taxonomic groups. Numbers show the average coverage these strains represent on the species and higher taxonomic levels in a collection of 219 microbiome samples from healthy individuals ^21,22^. For example, the 22 strains account on average for 59.6% of all bacterial biomass on a species level, 83.7% on a genus level, etc. (**B**) Breakdown of the sample-to-sample variation of this coverage. Coverage for additional studies is shown in the **Data S2**.

**Figure S3.**
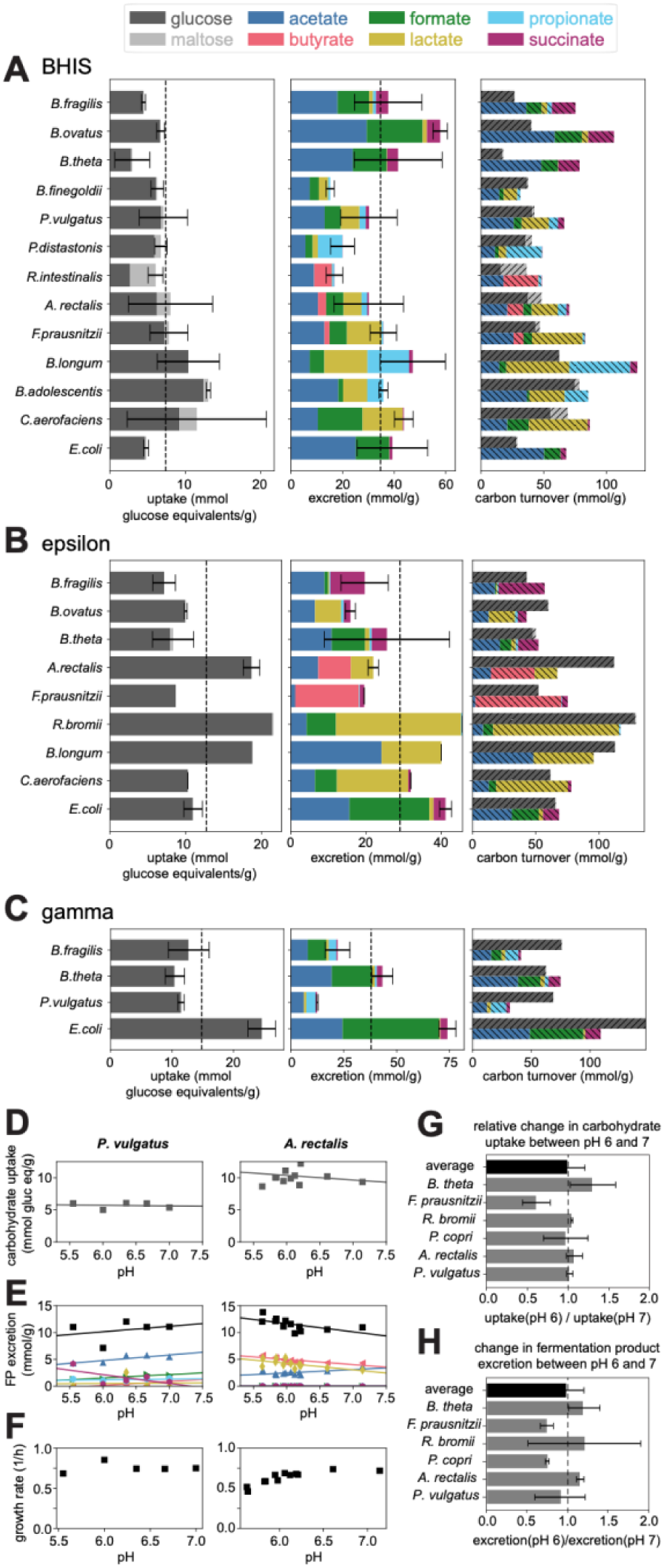
Uptake and excretion characteristics for growth in different conditions, Related to Figure 1. (**A**-**C**) Sugar uptake and fermentation product excretion per biomass for growth of different gut species in three different media (BHIS, 𝛾, and 𝜖). Complexity of medium composition decreases from BHIS (**A**) to YCA (Figure 1 D-F) to epsilon (**B**) and gamma medium (**C**). See **STAR Methods** for details on media composition. With decreasing medium complexity, fewer of the 22 strains can grow. Strain-by-strain visualizations of growth curves, fermentation product excretion, and sugar uptake for all media conditions and biological replicates can be found in the paper’s GitHub repository. (**D**-**H**) Sugar uptake and fermentation product excretion per biomass for growth at different pH values of 6 abundant bacterial species representing the most abundant genera and families. pH variation of carbohydrate uptake per dry mass (**D**), fermentation product excretion per dry mass (**E**), and growth rates (**F**) for *P. vulgatus* and *A. rectalis*, respectively. Liner trends with pH were fitted to determine the relative change in uptake (**G**) and excretion (**H**) between pH 6 and 7. Data for *B. thetaiotaomicron* and *A. rectalis* from^73^.

**Figure S4.**
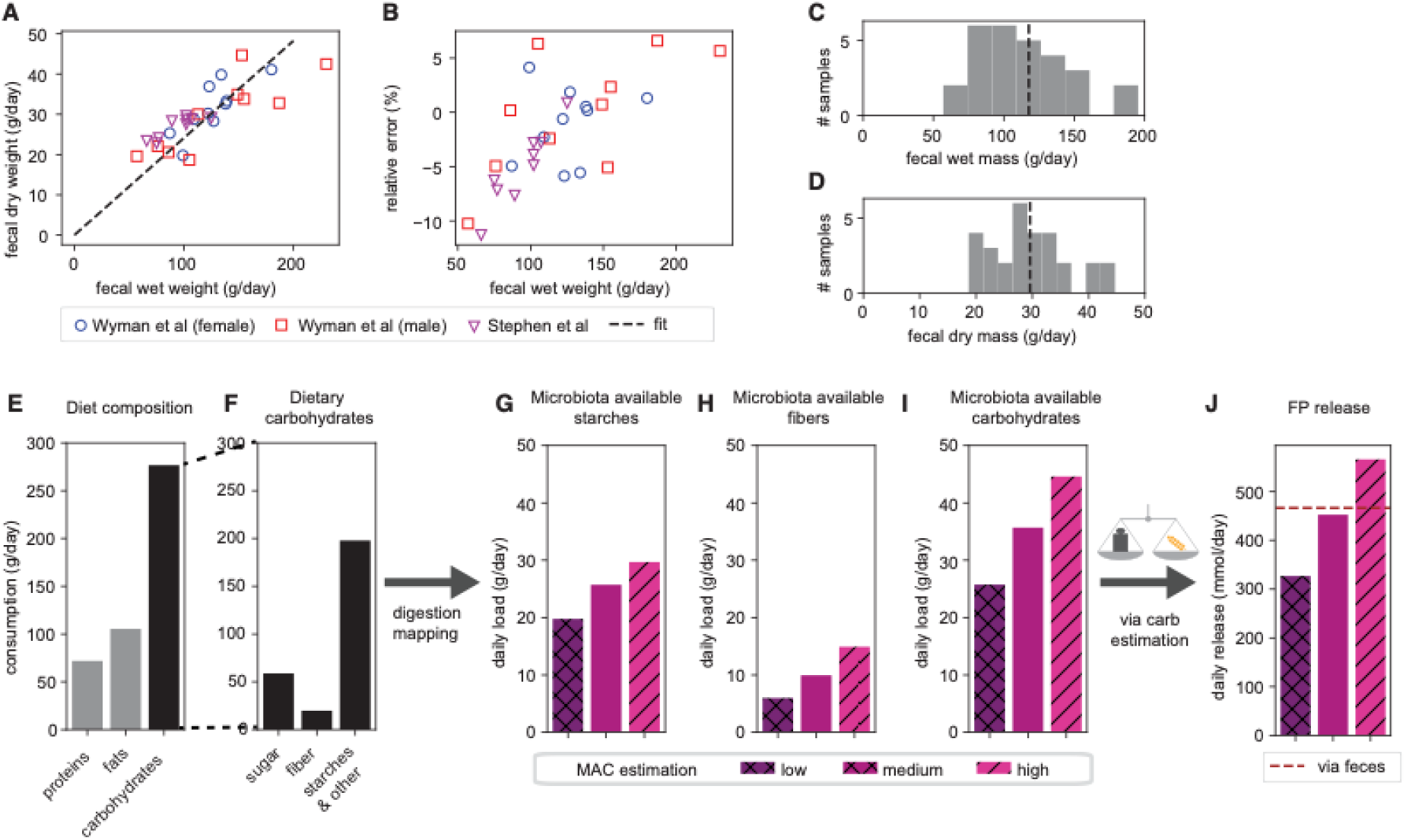
Characteristics of the British reference diet and variation of the microbiota available carbohydrates with different efficiencies of complex carbohydrate digestion, Related to Figure 2. To describe the relation between fecal dry and wet weight in people consuming a British reference diet, we use data from ^25^ where both values are reported. (**A**) linear regression (dashed line) assuming a fixed fraction of dry weight per wet weight describes the relation well. (**B**) Relative error of the linear regression model. Errors smaller than 10% confirm the linear model. (**C, D**) Histograms of fecal wet and dry weight to illustrate their substantial variation. (**E, F**) Diet composition of the British reference diet as reported in ^27^. (**G, H, I**) Digestibility of starches changes with the type of food consumed and the cooking method. The ability to digest different types of fibers also depends on the specific metabolic capabilities of the microbes present. To illustrate the effect of this variation, we use three different mappings from dietary carbohydrates to microbiota available carbohydrates (low, medium, high), accounting for the large variation in fiber digestibility (see **Supplementary Text Section 4**). These assumptions lead to varying amounts of microbiota available starches and fibers, the sum of which is the total amount of microbiota available carbohydrates. The medium case shows the best-established mapping for the British reference diet and is used in the main text. (**J**) The different mapping assumptions lead to different estimates of fermentation product release. Parameters used for MAC mapping follow typically observed ranges of starch content and fiber digestion, with 15% starch passage and 75% microbial fiber digestion for the high case (poor complex carbohydrate digestion along the upper digestive tract), 13% starch passage, and 50% fiber digestion for the medium case, and 10% starch passage and 30% fiber digestion for the low case (efficient complex carbohydrate digestion along the upper digestive tract).

**Figure S5.**
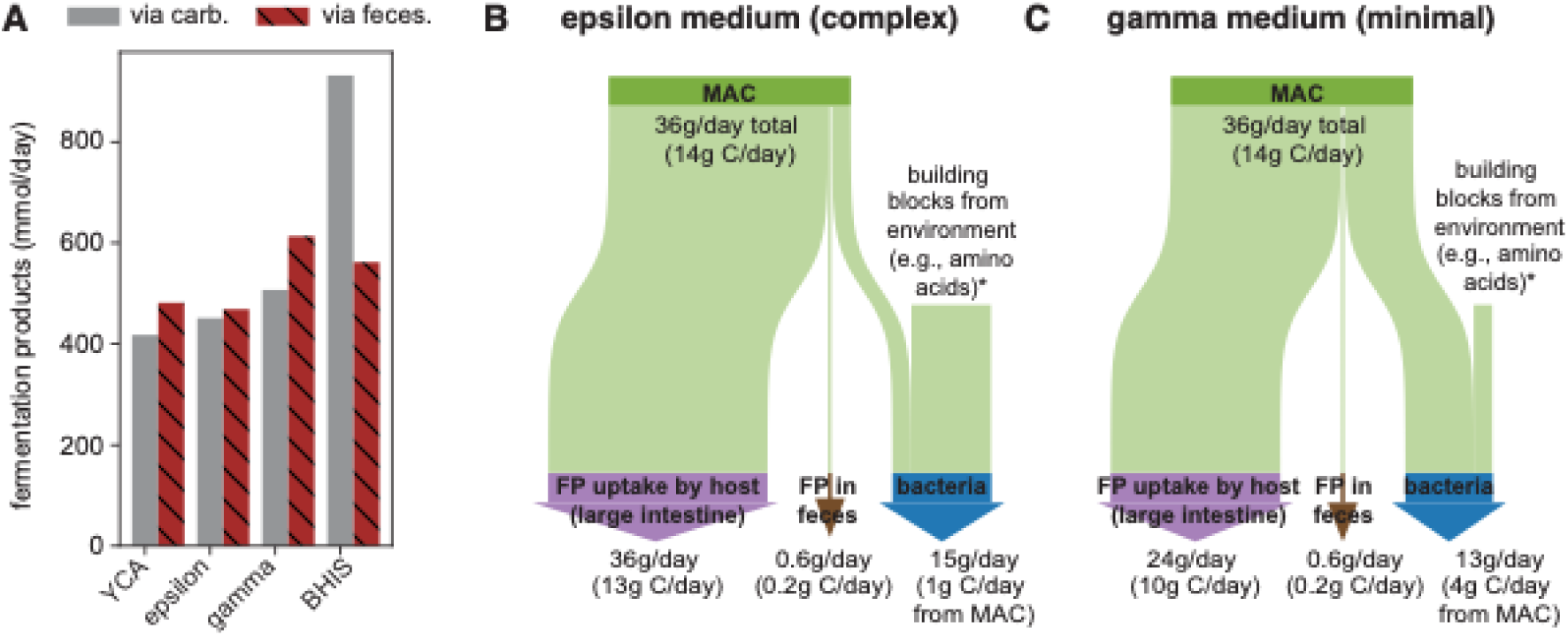
Variation in estimates of fermentation product harvest with media, Related to Figure 2. **(A)** Estimation via feces and carbohydrates for the British reference scenario, using average per-biomass uptake and excretion rates measured for the different media (**Figure S3**). The resulting daily release of fermentation products is comparable across media, except for the estimation via carbohydrates in BHIS. This is likely an artifact of the unknown carbon sources in this medium, the consumption of which we are unable to measure, leading to an underestimation of the per-biomass carbohydrate uptake rate. (**B,C**) Flow diagrams showing the resulting carbohydrate and carbon flow along the large intestine when using characteristics measured in gamma and epsilon media, respectively. Diagrams similar to the one shown in Figure 2E for YCA medium, but, as fewer strains grew in these media, we used simple rate averages and did not weight rates by biomass abundance. An internally consistent flow diagram for BHIS was not constructed due to the underestimation of carbohydrate intake in this medium.

**Figure S6.**
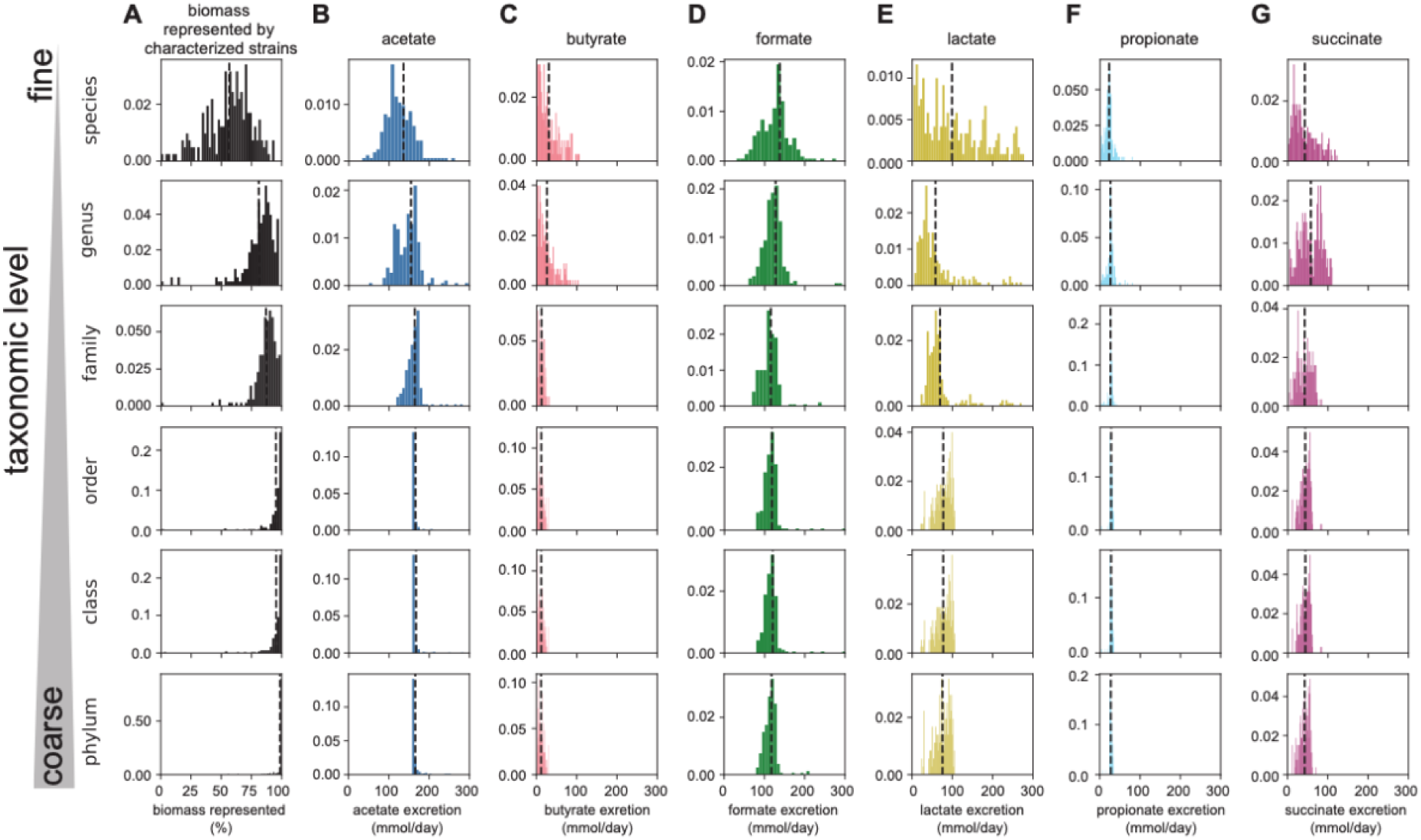
Representation and analysis of fermentation product release on different taxonomic levels, Related to Figure 3. To decide which taxonomic level to use for our analysis, we probed how much of the abundance in the healthy human gut microbiome ^21,22^ is represented by the 22 strains we have characterized experimentally. Covered abundance changed with the taxonomic level we chose for our analysis, with a coarser level leading to higher coverage (grey histograms). For example, on the species level, the characterized strains represent an average of about 60% of microbial biomass, while on the phylum level, the strains represent, on average, more than 90% of bacterial biomass. Thus, choosing a coarser level will lead to a higher representation by experimentally characterized strains, and we need to make fewer assumptions on how to describe the uptake and excretion rates of experimentally non-represented bacterial biomass. On the other hand, we also lose resolution in describing the production of specific fermentation products when choosing a taxonomic level that is too coarse. Particularly, the metabolic behaviors of strains belonging to the same taxonomic group might differ substantially at coarser taxonomic levels. For example, consider the production of succinate, which is produced in substantial quantities by some, but not all, species of the phylum Bacteroidota. When using average uptake and excretion rates, including all experimentally characterized strains belonging to the Bacteroidota phylum, we will lose possible differences in succinate release. As the best compromise, we thus chose the genus level for our analysis in the main text, as biomass coverage is already very high while we avoid too much averaging out of strain-level differences. See **Supplementary Text Section 2** for further discussion.

**Figure S7.**
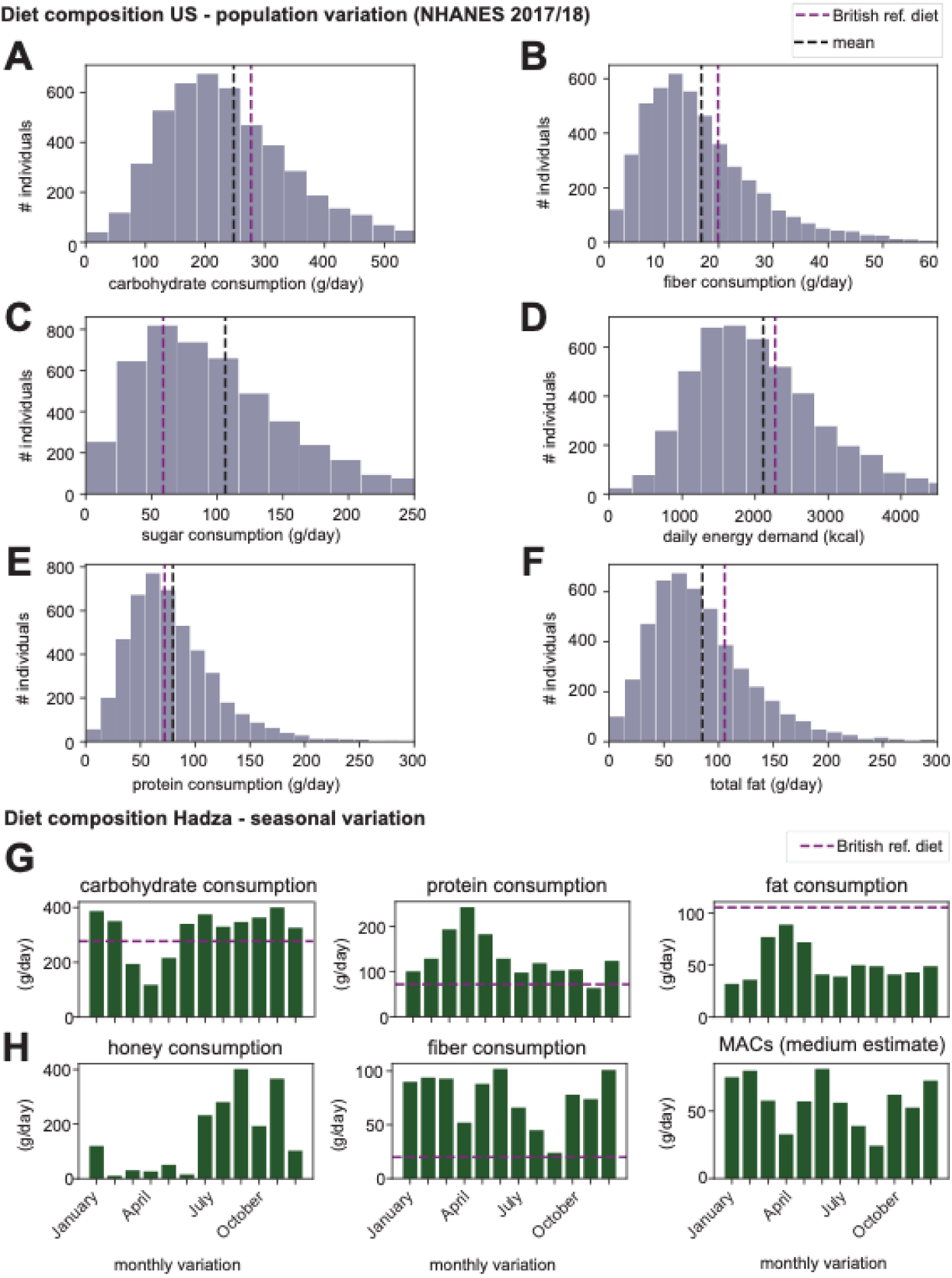
Major dietary components for the US and the Hadza populations, Related to Figure 4. This data underlies the estimates for microbiota available carbohydrates and total fermentation product release shown in Figure 3. (**A-F**) Distribution of different dietary characteristics across the US population based on the NHANES 2017/2018 cohort ^32^. Black dashed lines indicate means of distributions. (**G,H**) Per-capita breakdown of major diet components based on food collected by a group of Hadza people, as reported by Pontzer and Wood ^36^. Purple dashed lines in different panels indicate corresponding numbers for the British reference diet (**Figure S4**).

**Figure S8.**
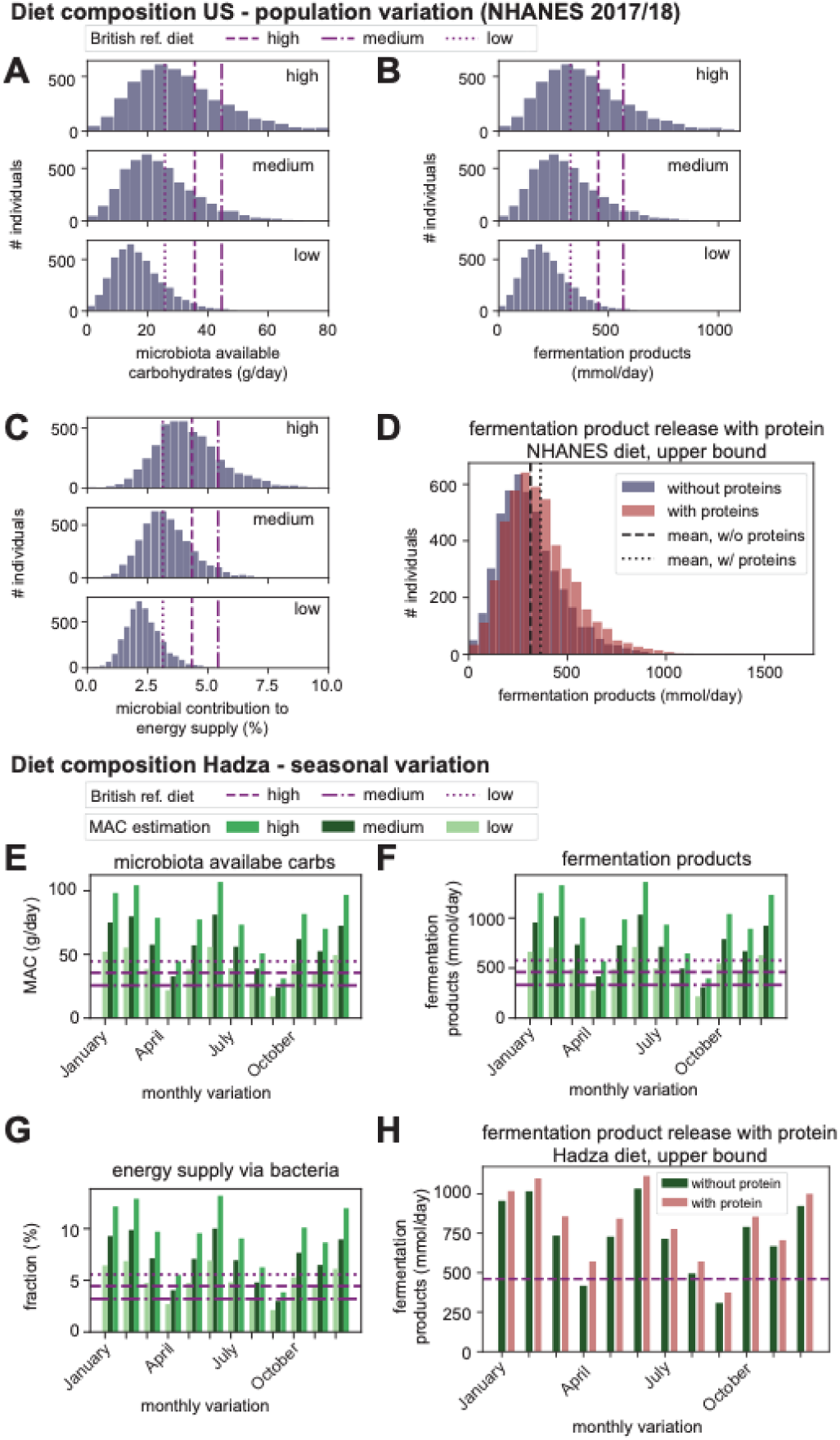
Variation of fermentation product release with diet, given different assumptions of carbohydrate digestion, Related to Figure 4. We analyzed the change in microbiota available carbohydrates for different diets to estimate the total fermentation product release when changing the assumptions on carbohydrate degradation along the upper digestive tract within observed ranges (**Figure S4**, **Supplementary Text Section 4** for further discussion of the mapping). (**A-C**) Changes in microbiota available carbohydrate, total fermentation product release, and microbial contribution to the daily energy demand with reported variations in carbohydrate consumption reported for the US NHANES 2017/2018 cohort. (**D-F**) Changes in microbiota available carbohydrates, total fermentation product release, and microbial contribution to the daily energy demand with the reported month-to-month variation of per-person carbohydrate amount in food in a group of Hadza people. (**G**) Fermentation product release for the NHANES cohort with (red) and without (blue) accounting for the fermentation of dietary proteins. (**H**) Fermentation product release for the Hadza people with (red) and without (green) accounting for the fermentation of dietary proteins. Changes in (G) and (H) are small, even for the used upper bound estimation of bacterial protein digestion (**Supplementary Text Section 5**). Following data for the British reference scenario (**Supplementary Text Section 4**), parameters used to describe the mapping between consumed and microbiota available carbohydrates are 15% starch passage and 75% fiber digestion for the high case, 13% starch passage and 50% fiber digestion for the medium case, and 10% starch passage and 30% fiber digestion for the low case. In the main text, data for the medium case are shown. Purple lines indicate corresponding estimations for the British reference diet.

**Figure S9.**
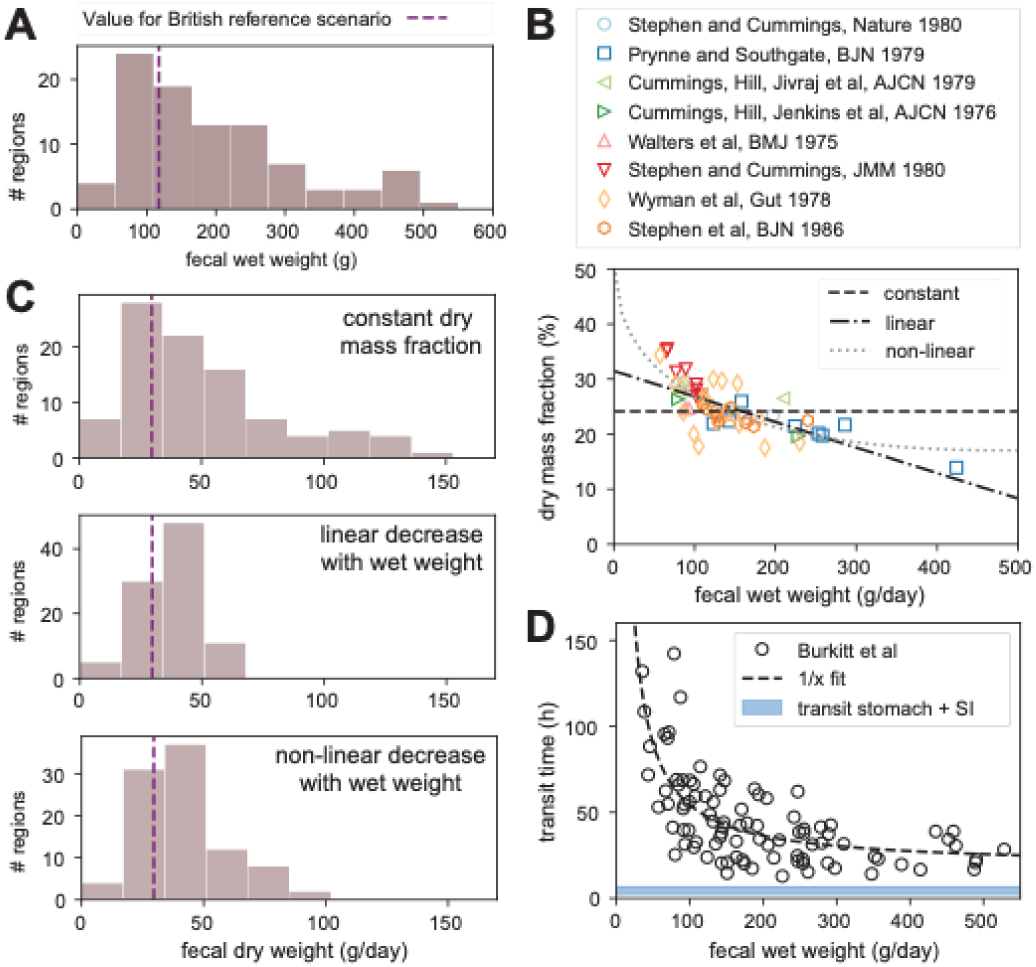
Global variation in fecal weight and variation of fermentation product release with fecal water content, Related to Figure 4. (**A**) Variation in fecal wet weight in humans of various backgrounds and lifestyles, as reported in ^38^. (**B**) Shown data (markers) were collected from different studies that measured fecal wet and dry weight for different cohorts and dietary compositions ^17,25,26,39–43^. When the variation in fecal output gets larger, water content in feces increases. Therefore, the assumption of a constant ratio of dry weight per wet weight we have been using for the British reference case (**Figure S4**) leads to substantial errors in describing the observations (“constant”, dashed line). To account for larger variations in water content, we formulated two additional models to fit the data. In the linear model, the dry mass fraction (𝛼_𝑑𝑤_) decreases linearly with fecal wet weight (“linear”, dotted dashed line, 𝛼_𝑑𝑤_ = 1 − 4.63 ∗ 𝑀_𝑓𝑒𝑐𝑒𝑠,𝑤𝑒𝑡_ + 0.69). In the non-linear model, the dry mass fraction (𝛼_𝑑𝑤_) decreases approximately linearly with fecal wet weight when wet weight is low, but it hardly changes anymore when fecal wet weight is high (“non-linear”, dotted line, 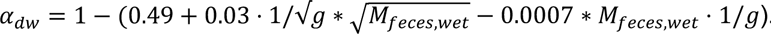. Both models describe the trends of the available data well. We expect the dry mass content to decrease further with higher fecal wet weight as very high weights are likely caused by a higher water content bound to fibers in feces. However, as no parallel measurements of fecal dry and wet weights are available for very high weights, the non-linear model provides a well-supported upper bound of fecal dry weight. (**C**) With these different models, we then estimated the fecal dry weight for the data reported in ^38^. Notably, the variation in fecal dry weight decreases substantially when accounting for the adjustment in dry mass fraction. For the estimations discussed in the main text, we used the non-linear model. As such, these numbers provide an upper bound estimate. (**D**) The transit time (i.e., the time it takes for ingested material to pass the entire digestive tract), decreases with fecal wet weight, described well by an inversely proportional relation (dashed line, 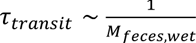). Data re-plotted from Burkitt et al. ^38^. As transit through the stomach and small intestine is relatively short (highlighted in blue), this strong variation in transit time is primarily due to changes in the time biomass remains within the large intestine. Notably, the substantial variation of this time can impact the efficiency with which the microbiota digests complex carbohydrates along the large intestine. Particularly when fecal wet weight is high and transit times are short, the microbiota might simply not have enough time to complete digestion, and a higher fraction of complex carbohydrates end up undigested in feces. As such, and similar to the estimation of fecal dry weight, our estimations provide again an upper bound for the daily production of fermentation products, particularly when fecal wet weights are large.

**Data S1. Variation of fermentation product release with microbiome composition and diet, Related to Figures 3 and 4**. This interactive figure (available as supplementary html file or via https://cremerlab.github.io/fermentation_products/study-explorer) facilitates the analysis of fermentation product release with varying microbiome composition and diet composition utilizing our analysis framework (**Supplementary Text Section 2 and 3**). Adjustment of the figure settings works in two steps. First, microbiome composition can be selected from a collection of metagenomics studies^31^ which includes data from about 83.000 fecal samples (top panel). The selection menus allow for the specification of studies, age category, and health status. The grey histogram (right) illustrates what fraction of the microbial biomass is represented across samples by the bacterial species we experimentally examined in this study. Second, diet dependent parameters can be chosen (middle panel), either by setting directly the amount of microbiota available carbohydrates, by adjusting the amount of dietary starch and fibers consumed, or by selecting a preset diet introduced in the main text (British reference diet, mean of NHANES US diets, or mean of Hadza diets). Based on these selections of microbiome samples and dietary parameters the distributions of single and total fermentation products are shown (bottom panel). Additionally, we show the total daily energy content of all secreted fermentation products. The analysis, including the characterized biomass and release of different fermentation products, is based on the genus level with experimentally characterized species representing the genus (**Supplementary Text Section 2 and 3**). Diet mapping as described in **Supplementary Text Section 4**. A similar analysis based on new metagenomics data or dietary information can be realized using the scripts provided on the GitHub repository. The code to generate the interactive figure is also available via the GitHub repository.

**Table S1. Per biomass uptake and excretion of metabolites, and their molecular properties, Related to Figure 1**. Based on in-vitro measurements of all 22 strains as outlined in **Supplementary Text Section 2 and Figure S1**. Data for growth in YCA medium. Unweighted averages and ranges observed across all strains. In the calculations of fermentation product and energy harvest, we additionally account for the relative abundance of the different strains (**Supplementary Text Section 2**). **\*** Enthalpy values based on^74^. ** While formic acid (formate) is an abundant fermentation product that can enter the host’s circulation system, there is no evidence that it contributes directly to energy generation by the host. Therefore, we do not include formic acid when estimating the fermentation product contribution to host energy demand.

## STAR Methods

### KEY RESOURCES TABLE

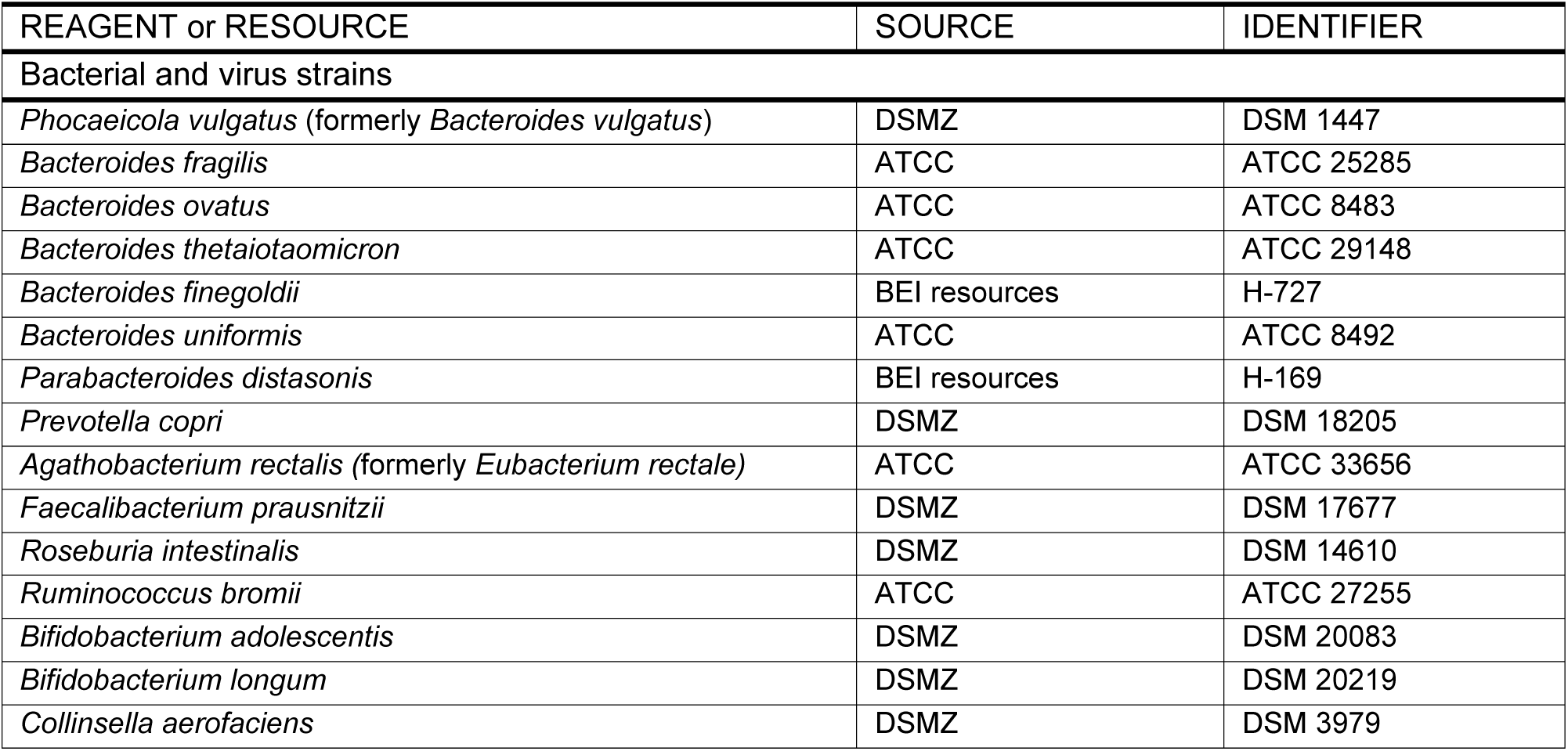
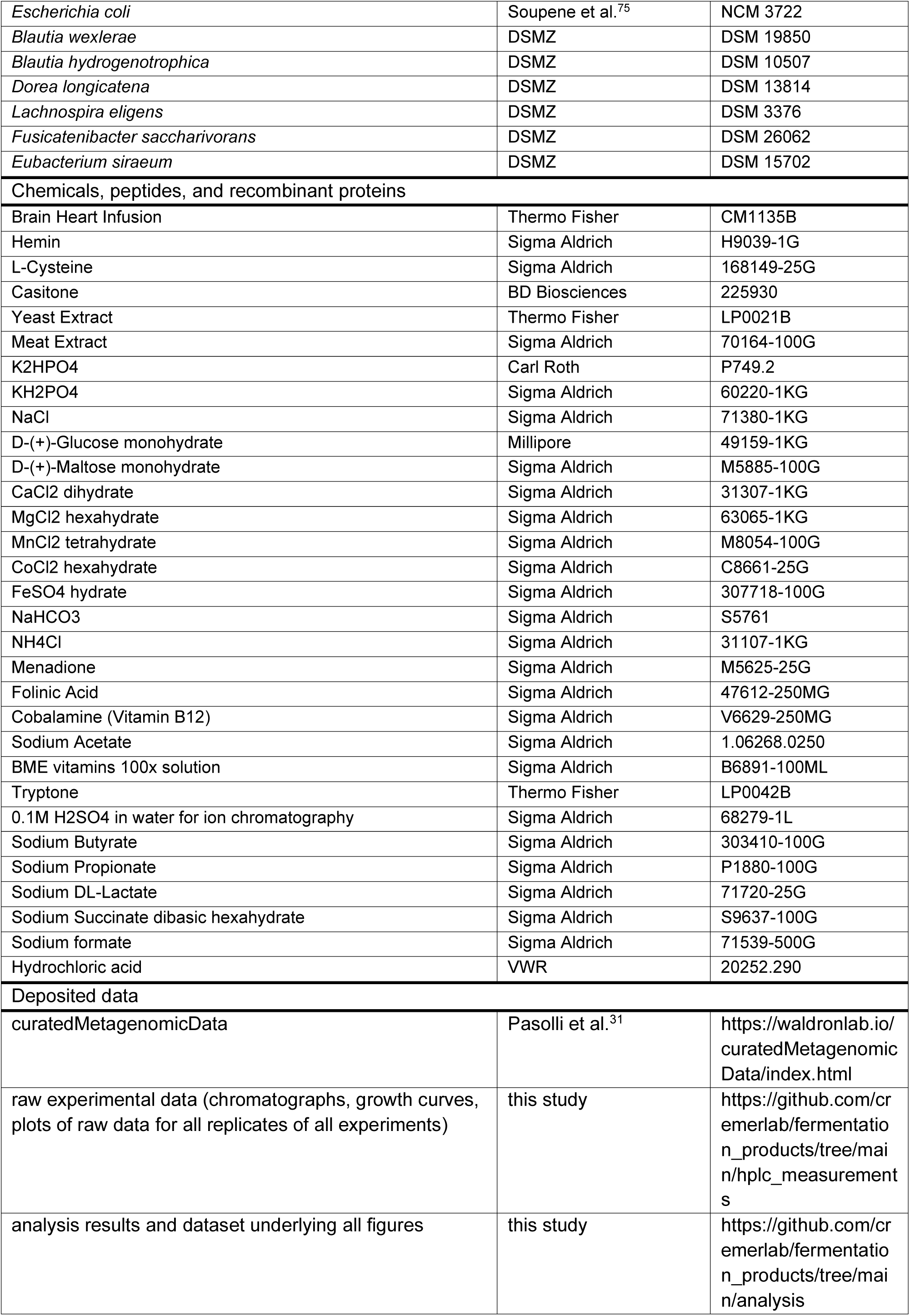
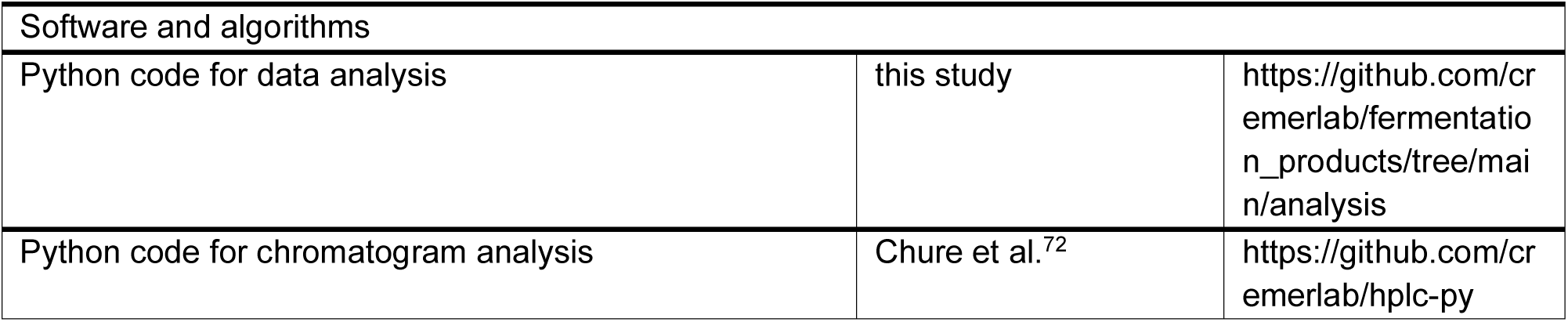

### EXPERIMENTAL MODEL AND STUDY PARTICIPANT DETAILS

#### Bacterial strains and culturing conditions

All characterized strains were obtained from established strain repositories and are listed in the **Key Resources Table**, along with details for media components and chemicals used. Species identity of all strains was confirmed by 16S rDNA sequencing (PCR amplification followed with Sanger sequencing). Bacteria were grown in the media indicated in the respective figures. Media used were of different complexities, all include ammonia as nitrogen source and glucose or maltose as carbon sources. BHIS medium is Brain Heart Infusion supplemented with 10mg/L hemin (prepared as in^76^) and 1g/L L-cysteine. YCA medium is a modified version of YCFA^77^, containing 1% casitone, 0.25% yeast extract, 0.1% meat extract, 4g/L NaHCO_3_, 1g/L cysteine, 0.45g/L K_2_HPO_4_, 0.45g/L KH_2_PO_4_, 0.9g/L NaCl, 0.09g/l MgSO_4_•7 H_2_O, 0.12 g/L CaCl_2_•2H_2_O, 10mg/L hemin, 0.02mg/L cobalamin, 0.06mg/L p-aminobenzoic acid, 2mL/L BME vitamins, 10mM sodium acetate, 20mM NH_4_Cl, and 20mM glucose or 10mM maltose, respectively. For the data shown in **Figure S4** (except for the data shown for *B. thetaiotaomicron* and *E. rectale*; these have been published in^14^, and they were grown in epsilon medium as described there), YCA was used as described above, but the phosphate buffer concentration was increased to 100mM in the final medium. pH was adjusted by changing K_2_HPO_4_/KH_2_PO_4_ ratios, and for low pH HCl was added to achieve the desired pH, which was checked using a benchtop pH meter (Mettler Toledo SevenCompact pH with InLab Routine electrode). Gamma medium is a fully defined medium, containing 72mM K_2_HPO_4_, 28mM KH_2_PO_4_, 50mM NaCl, 0.5mM CaCl_2_, 0.4mM MgCl_2_, 0.05mM MnCl_2_, 0.05mM CoCl_2_, 0.004mM FeSO_4_, 5mM Cysteine, 20mM NaHCO_3_, 20mM NH_4_Cl, 1.2mg/mL Hemin, 1mg/mL menadione, 2mg/mL folinic acid, 2mg/mL cobalamine, 10mM acetate, 0.2% BME vitamins, and 20mM glucose. Epsilon medium has the same composition as gamma medium, but also contains 1% Tryptone. All bacteria were cultured in glass tubes in anaerobic work benches (Coy Labs), in temperature controlled dry baths (Eppendorf ThermoMixer C) at 37°C, shaking with 450rpm. Cultures were inoculated from glycerol stocks stored at −80°C, which were transferred to the anaerobic work benches before opening. Except where explicitly stated, experiments were performed at neutral pH values.

### METHOD DETAILS

#### Growth experiments and sampling for metabolite analysis

For growth experiments, glycerol stocks of bacterial cells were taken from a −80°C freezer to an anaerobic workbench. Cells were then transferred from stocks to sterile glass tubes with YCA or BHIS media (seeding culture). After substantial growth, as indicated by an optical density at 600nm (𝑂𝐷_600_) over 0.1, cultures were diluted into new culture tubes with YCA (pre-culture). Before cells reached saturation, 𝑂𝐷_600_ < 1, cultures were then diluted again into fresh media to 𝑂𝐷_600_ ≈ 0.02 to start the experimental cultures. Growth was then tracked by regular measurements of 𝑂𝐷_600_ (around 7 measurements between 𝑂𝐷_600_ 0.05 and 0.5). 𝑂𝐷_600_ measurements were performed in Quartz semi-micro-cuvettes in a benchtop spectrophotometer (Thermo Scientific Lambda/Genesys30) or by directly determining optical density of the culture tubes. We also collected four to six samples of 200µl culture volume for the subsequent quantification of sugar and fermentation products concentrations. These samples were transferred to sterile 0.2µm filter centrifuge tubes (Corning Costar Spin-X centrifuge filter tubes) and centrifuged at 11000g for 2min to remove cells. Samples were then kept at 4°C until further analysis. For all samples, optical densities were typically kept between 𝑂𝐷_600_ ≈ 0.04 to 0.5 to ensure linearity between absorbance and bacterial biomass and avoid substantial accumulation of metabolites (such as fermentation products) that could affect growth conditions. To ensure pH stability, we regularly measured pH at the end of growth experiments (𝑂𝐷_600_ ≈ 0.5). All measurements were done in YCA, BHIS, epsilon, or gamma media, typically for three biological replicates each. Not all strains could grow in all media, and we report only results for strains with substantial growth (𝑂𝐷_600_ ≳ 0.1) within 48h of inoculation in the respective media. Media compositions are provided above.

#### Chromatography Method to Determine Nutrient Uptake and Fermentation Product Secretion

Metabolite concentrations in filtered samples from culture supernatants were analyzed using isocratic HPLC with refractive index detection (Thermo Scientific Ultimate 3000/Shimadzu Prominence SC2030C), as described in^14^. In short, 20µl of sample was injected using an autosampler cooled to 10°C, separated over an ion exchange column (Phenomenex Rezex RoA organic acid H+ (8%), LC column 300 x 7.8mm) in a column oven at 40°C, at a flow rate of 0.4ml/min, with 2.5mM H_2_SO_4_ in water as mobile phase. Chromatography data from the RID detector was recorded for 40min, exported as plain text files, and analyzed using custom Python scripts which have subsequently been released as a standalone software package^72^. An example of the analysis is discussed in **Figure S1** and the analysis scripts are available on the GitHub repository of this study. In short, a background correction based on the algorithm described in^78^ was applied and areas under all the peaks representing the metabolites of interest were extracted. These areas were compared to a standard curve of known metabolite concentrations, and values for uptake or excretion per biomass were then calculated for every growth experiment.

### QUANTIFICATION AND STATISTICAL ANALYSIS

#### Estimating daily fermentation product release

To calculate the daily release of fermentation bacteria in the large intestine we started with estimations on the bacterial biomass released daily in feces 𝑀_𝑓𝑒𝑐,𝑏𝑎𝑐_, or the daily amount of complex carbohydrates which reach the large intestine and are digestible by bacteria, 𝑀_𝑐𝑎𝑟𝑏_. These values either follow from fecal mass quantifications^25^ and the characterization of fecal matter^26^, or from a mapping estimating the link between consumed carbohydrates and those available for the microbiota to utilize (**Supplementary Text Section 4**). We then calculated the daily release as 𝐹𝑃_𝑡𝑜𝑡_ = 𝜖_𝑡𝑜𝑡_ ⋅ 𝑀_𝑏𝑎𝑐𝑡_ or 𝐹𝑃_𝑡𝑜𝑡_ = 𝜖_𝑡𝑜𝑡_ ⋅ 𝑌_𝑐𝑎𝑟𝑏_ ⋅ 𝑀_𝑐𝑎𝑟𝑏_ (**Supplementary Text Section 3**) with the total per biomass excretion 𝜖_𝑡𝑜𝑡_ and Yield values 𝑌_𝑐𝑎𝑟𝑏_ following from the values of experimentally characterized strains, weighted by the biomass abundance these strains represent on the genus level. The weighting approach is described in detail in **Supplementary Text Section 2**. To illustrate the variation of fermentation product harvest with microbiome composition (**Figure 3**), we weighted the uptake and excretion values of different strains based on their relative abundance in 219 different samples from healthy adults^22^. For other harvest estimations in the main text (**Figures 2 and 4**), we used average abundance numbers observed across the same 219 samples. In addition, we provide an interactive figure (**Data S1**) allowing the application of our framework to the database of metagenomic studies available in the Curated Metagenomics Dataset ^31^.

#### Calculation of variation with diet

To obtain variations with diet, we analyzed different datasets on dietary composition and fecal weight as introduced. From dietary data we estimated the variation in microbiota available carbohydrates which we then used to calculate the variation in the daily release of fermentation products using the via-carbohydrate estimation (**Figures 2BC and S8, Supplementary Text Section 3**). From fecal wet weight data we estimated the variation of daily bacterial biomass released in feces, accounting for changes in fecal water uptake (**Figure S9**). With this data we then calculated the daily fermentation product release using the via-feces estimation (**Figure 2A and Supplementary Text Section 3**).

#### Statistical analysis and error propagation

Where not stated otherwise, we used N=3 biological replicates to determine fermentation product excretion and carbon source uptake. Error bars in figures indicate standard deviations of those replicates (**Figure 1, Figure S3**). To estimate variation in our quantitative analysis framework, we used Gaussian error propagation based on standard deviations of the underlying data (**Figures 2, 5, 6**).

### ADDITIONAL RESOURCES

Our customized Python scripts, raw experimental data including chromatography data and growth curves), as well as processed data, are available on the paper’s GitHub repository (https://doi.org/10.5281/zenodo.10445504) accessible via https://github.com/cremerlab/fermentation_products. The code to generate the interactive figure (**Data S1**) is available via the GitHub repository, and the interactive figure can be accessed at https://cremerlab.github.io/fermentation_products or by opening the supplementary HTML file.

