## Supplementary Text for "Quantifying the varying harvest of fermentation products from the human gut microbiota"

#### Table of Contents

1. ANAEROBIC GROWTH IS CONSTRAINED BY BALANCING ATP GENERATION AND REDOX BALANCE 2
1.1. Energy generation and redox balance in different metabolic pathways 2
1.2. ATP yields and fermentation product release per biomass across media and ecological conditions 4
2. MEASURING CARBOHYDRATE UPTAKE AND FERMENTATION PRODUCT EXCRETION 5
2.1. Uptake and excretion values from measured data 5
2.2. Calculating averages weighted for abundance in reference microbiomes 6
3. CALCULATION OF FERMENTATION PRODUCT RELEASE 8
3.1. Estimation via feces 8
3.2. Estimation via carbohydrates 9
3.3. Estimation via theoretical ATP yields 9
4. MAPPING OF NUTRITION DATA TO MICROBIOTA-AVAILABLE CARBOHYDRATE 10
5. CARBON SOURCES NOT DERIVED FROM DIETARY CARBOHYDRATES 12
5.1. Protein-derived fermentation products 12
5.2. Mucin-derived fermentation products 13
5.3. Contribution of protein and mucin fermentation to overall release of fermentation products 15
6. MOST RELEASED FERMENTATION PRODUCTS ARE TAKEN UP BY THE HOST 15
7. MOUSE ESTIMATIONS 16
7.1. Estimation via feces 16
7.2. Estimating microbial energy output by comparing energy retention in germ-free and colonized mice 17
7.3. Estimating relevant amounts of fermentation products in humans and mice 18
8. ESTIMATION OF ENERGY CONTRIBUTION 19
9. REFERENCES 19

### 1. Anaerobic growth is constrained by balancing ATP generation and redox balance

#### 1.1. Energy generation and redox balance in different metabolic pathways

Bacteria growing in anoxic conditions face a threefold challenge: they need to generate sufficient energy, retain enough carbon to produce biomass, and keep their redox state in balance [1]. In this section, we briefly summarize the biochemical origin of this challenge for gut bacteria. To survive and grow, bacteria need to extract energy and building blocks for biomass synthesis from nutrients available in their environments. Under anoxic conditions, bacteria also need to find a suitable terminal electron acceptor instead of oxygen, making anaerobic metabolism much less energy efficient than aerobic growth in the presence of oxygen. Biochemically, this emerges from the fact that most energy-providing metabolic reactions are redox processes. The electron-donating and electron-accepting parts of these reactions are typically separated and coupled by electron carriers (often nicotinamide adenine dinucleotides, NAD). The donating reaction transfers electrons to the carrier, the accepting reaction transfers electrons from the carrier to an accepting molecule, and both reactions can be coupled with the generation of ATP. Importantly, cells need to balance their redox state, for which every electron-donating reaction needs to be coupled with an electron-accepting reaction. Most obligate anaerobes within the gut use metabolic intermediates as terminal electron acceptors, which are then secreted as fermentation products. These compounds, typically small organic molecules, contain carbon extracted from nutrients that therefore cannot be used to gain more energy or build biomass.

In the large intestine, the main carbon source that is available to the microbiota are carbohydrates that are undigestible by host enzymes (we also discuss the role of proteins and mucin in **Supplementary Text Section 5** below). After bacterial digestion, carbohydrates get metabolized to pyruvate via glycolysis or similar pathways and are then funneled into a limited set of pathways leading to the final fermentation products (**Figure 1A**). Not all bacterial species carry all these pathways, and every given bacterial strain faces the optimization problem discussed above: maximizing growth and survival while keeping its metabolism, including ATP concentrations and redox states, in homeostasis. Importantly, however, these pathways are part of the central carbon metabolism, and even though there is variation between species and sometimes even between strains of the same species in pathway utilization, the metabolic abilities are typically conserved within genera and often families (see e.g. [2,3] for *Bacteroides* strains, [4,5] for analyses of the phylogenetic distribution of propionate production pathways, [5–7] for analyses of the phylogenetic distribution of butyrate production pathways). We have analyzed how the choice of taxonomic level influences our results (**Supplementary Text Section 2** below and **Figure S6**) and chose the genus level as taxonomic level discussed in the main text.

In the following, we summarize the most abundant fermentative pathways that are found in gut bacteria from the different families and briefly mention additional details. Importantly, the shown net reactions are one possible outcome that preserves stoichiometry, but other combinations are possible.

**Bacteroidetes** (simplified, [8–10]): this combination of pathways is typical for members of the Bacteroidaceae family, represented by *P. vulgatus*, *B. fragilis*, *B. ovatus*, *B. thetaiotaomicron*, *B.*

*finegoldii*, *B. uniformis* in our dataset. Prevotellaceae (represented by *P. copri*) have very similar pathways, but lack the last step for propionate formation and secrete succinate instead [11]. Also members of the Bacteroidaceae family can skip this last step of the propionate formation pathway and secrete succinate, which has been suggested to be subject to regulation [12].

1. 1 Glucose + 2 $NAD^+$ → 2 Phospoenolpyruvate (PEP) + 2 NADH
2. 2 PEP + 3 ADP + 2 NADH → 1 Pyruvate + 1 Propionate (or Succinate) + 3 ATP + 2 $NAD^+$
3. 1 Pyruvate + 1 ADP → 1 Acetate + 1 ATP

(4. 1 Pyruvate + NADH → 1 Lactate + 1$NAD^+$)

(5. 1 Pyruvate → 1 Formate)

**Net reaction: 1 Glucose + 4 ADP → 1 Propionate (or Succinate) + 1 Acetate + 4 ATP**

**Clostridia** (simplified, [6,8,13]): this combination of pathways is typical for abundant gut dwelling members of the class Clostridia, represented by *A. rectalis*, *R. intestinalis*, and *F. prausnitzii* in our dataset. For *A. rectalis* and *R. intestinalis*, we observe net production of acetate, which is not covered in the net reaction shown below.

1. 1 Glucose + 2 NAD+ + 2 ADP → 2 Pyruvate + 2 NADH + 2 ATP

2a. 1 Pyruvate + 1 NADH → 1 Lactate + 1 $NAD^+$

2b. 1 Pyruvate + 1 ADP → 1 Acetate + 1 ATP

3. 1 Acetate + 1 ATP + 1 CoA → 1 Acetyl-CoA + ADP
4. 1 Acetate + 1 NADH + 1 ADP + 1 Acetyl-CoA → 1 Butyrate + 1 ATP + 1 $NAD^+$

**Net reaction: 1 glucose + 1 ADP → 1 Lactate + 1/2 Butyrate + 1 ATP**

**Bifidobacteria** [14]: this pathway is characteristic for members of the genus Bifidobacterium, represented by *B. adolescentis* and *B. longum* in our dataset.

1. 2 Glucose + 1 ADP → 2 Xylulose-5-P + 1 Acetate + ATP
2. 2 X-5-P + 6 ADP + 2 NAD+ → 2 Acetate + 2 ATP + 2 Pyruvate + 4 ATP + 2 NADH
3. 2 Pyruvate + 2 NADH → 1 Lactate + 1 NAD+ + 1 Formate + 0.5 Acetate + 0.5 ATP + 0.5 Ethanol + $NAD^+$

**Net reaction: 2 glucose + 7.5 ADP → 3.5 acetate + 0.5 ethanol + 1 lactate + 1 formate + 7.5 ATP**

**Mixed acid fermentation** [15]: this combination of pathways is used by members of the Enterobacteriaceae family such as *Escherichia coli* and *Salmonella enterica* when growing in anaerobic conditions. In our dataset it is represented by *E. coli*. Ethanol, acetone, butanediol and butyrate have also been shown to be possible fermentation products, which we do not observe in our measurements.

1. 2 Glucose + 4 $NAD^+$ + 4 ADP $\rightarrow$ 4 Pyruvate + 4 NADH + 4 ATP
2. 1 Pyruvate + 1 ADP $\rightarrow$ 1 Acetate + 1 ATP
3. 1 Pyruvate + 1 NADH $\rightarrow$ 1 Lactate + 1 $NAD^+$
4. 1 Pyruvate + 1 ATP + 2 NADH $\rightarrow$ 1 Succinate (or 1 Propionate) + 1 ADP + 2 $NAD^+$

(5. 1 Pyruvate $\rightarrow$ 1 Formate)

**Net reaction: 2 Glucose + 6 ADP + 4 NADH $\rightarrow$ 1 Acetate + 2 Lactate + 1 Succinate + 6 ATP + 4 $NAD^+$**

**Acetogenesis** [8,16]: this pathway is used by rare members of the human gut microbiota, an is represented by Blautia hydrogenotrophica in our dataset.

1. Glucose + 2 $NAD^+$ + 2 ADP $\rightarrow$ 2 Pyruvate + 2 NADH + 2 ATP
2. 2 Pyruvate + 2 ADP $\rightarrow$ 2 Acetate + 2 ATP + 2 $CO_2$
3. $CO_2$ + NADH $\rightarrow$ Formate + NAD+
4. Formate + CO2 + ATP + NADH $\rightarrow$ Acetyl-CoA + ADP + NAD+
5. Acetyl-CoA + ADP $\rightarrow$ Acetate + ATP

**Net reaction: Glucose + 2 ADP + 2 NADH $\rightarrow$ 3 Acetate + 2 ATP + 2 $NAD^+$**

For completeness, we note that, in addition to substrate-level phosphorylation, anaerobic respiration can allow anaerobes to harvest energy from reactions that are insufficiently exergonic to produce ATP, e.g., via the formation of a chemiosmotic gradient coupled to an ATPase analogous to aerobic respiration. In the listing above, we did not include this possibility. However, in the phenomenological data shown in **Figure 1**, where we measure actual physiological parameters experimentally, we expect both classical fermentation and anaerobic respiration to be involved.

#### 1.2. ATP YIELDS AND FERMENTATION PRODUCT RELEASE PER BIOMASS ACROSS MEDIA AND ECOLOGICAL CONDITIONS

In the real gut, growth conditions vary strongly with ecological interactions in spatially heterogeneous niches that are likely impossible to replicate in a test tube [17,18]. The various conditions can drastically shape growth behavior and, thus, the detailed population dynamics that shape microbiota composition [19,20]. This strong variation in growth is, for example, illustrated by the growth rates we measured in pure cultures which strongly vary with media composition (see plots provided on data repository, folder hplc_measurements). However, despite this variation in growth behavior, the metabolic demands to support the synthesis of a specific amount of biomass are much more constrained and expected to vary far less with specific conditions. To be more specific, consider the energy requirement for biomass formation, by far the most energy-demanding process in growing bacteria [21,22]. To grow, bacteria need to synthesize novel biomass which requires energy, commonly quantified in the required amount of ATP per newly produced biomass. The ATP requirement to synthesize and assemble all cellular components has been well mapped out theoretically in thoroughly investigated model organisms, particularly for *Escherichia coli*. Notably, the theoretical ATP requirement varies little with the growth conditions cells encounter. For example, Stouthamer [23] reports that ATP requirements are almost identical when comparing growth in minimal media and a medium where amino acids are provided (34.7 and 34.9 mmol ATP per g dry biomass). This similarity is the result of very similar overall energy requirements for peptide elongation and amino acid synthesis / nitrogen uptake (minimal media) or amino acid uptake (amino acid supplemented media) needed to sustain peptide elongation. In the anaerobic environment of the gut, bacteria need to generate this energy via fermentation. As described above (**Supplementary Text Section 1.1)**, under these conditions ATP synthesis is coupled to the excretion of fermentation products to simultaneously enable energy supply and maintain redox balance. Furthermore, there is a limited number of fermentation pathways that gut bacteria use. In these pathways, typically between one and two molecules of ATP are synthesized per release of one fermentation product (**Supplementary Text Section 1.1**, not including formate). Accordingly, to generate the approximately 35mmol ATP theoretically required to synthesize one gram of bacterial biomass, bacteria would have to excrete 17 to 35 mmol of fermentation products. Notably, this theoretically predicted range recapitulates the observed excretion of fermentation products (**Figures 1**). Furthermore, the small change of theoretically required ATP amounts with media conditions rationalizes why we see so little change in excretion across media conditions (**Figures 1G and S3A-C**), further supporting our approach to infer the production of fermentation products within the human gut: even though it might never be possible to adequately recapitulate the ecological conditions in the gut in controlled laboratory conditions, the main metabolic parameters to sustain novel biomass synthesis in the gut are unlikely to differ dramatically from the values we measure in the laboratory. This expectation is further confirmed by our comparative analysis of fermentation product harvest and energy homeostasis in conventional and germ-free mice (**Supplementary Text Section 7**).

### 2. Measuring carbohydrate uptake and fermentation product excretion

#### 2.1. Uptake and excretion values from measured data

To estimate the daily amount of fermentation products released by the gut microbiota we experimentally characterized $N_{sp} = 22$ highly abundant species. Following the experimental workflow described in **Figure S1**, we characterized for each species $j$:

i. the molar amount of carbohydrates bacterial cells consume to support a given amount of bacterial biomass $\{u_j\}$. We denote this as the *per biomass uptake* and it follows from the sum of the experimentally determined glucose and maltose uptake ($u_j = u_{glucose,j} + 2u_{maltose,j}$, with mmol glucose equivalents as unit). The inverse of this value is commonly also denoted as yield coefficient ($Y_j = 1/u_j$).
ii. the excretion of major fermentation products per bacterial biomass, $\{e_{ij}\}$ with i = {acetate, butyrate, formate, lactate, succinate, propionate}, denoting the major fermentation products released and measured.

The variation of these numbers across strains is summarized in **Figure 1** and all values are listed in the data tables on the GitHub repository.

#### 2.2. Calculating averages weighted for abundance in reference microbiomes

To calculate average uptake and excretion rates in the gut from these measurements, we weighted measured rates by the abundance of different species in a microbiota sample. For example, if *B. uniformis* is the most abundant species in a sample, we also give the measured excretion behavior of this strain a higher weight when determining average excretion rates. We further used measured uptake and excretion rates of characterized strains to parametrize taxonomically closely related but uncharacterized strains, such as those from the same genus. The calculation is detailed in the following.

We started with the relative abundance $\alpha_j^{species}$ of each experimentally characterized species $j$. To obtain these abundance numbers for a large pool of microbiota samples, we use a large collection of metagenomics data (93 studies and about 18,000 samples from donors with different health conditions)[24]. In the simplest calculation, we use these abundance numbers to estimate the weighted average excretion per biomass for each fermentation product $i$:

$$e_i = \sum_{j \in charaterized\ strains} e_{ij} \cdot \tilde{\alpha}_j^{species}$$

Here, $e_{ij}$ denotes the measured excretion of fermentation product $i$ per biomass of characterized strain $j$. $\tilde{\alpha}_j^{species}$ is the abundance of species j, normalized by the abundance of all characterized species, $\tilde{\alpha}_j^{species} = \alpha_j^{species} / \sum_j \alpha_j^{species}$. We can similarly determine the uptake of carbohydrates per biomass as $u = \sum_j u_j \cdot \tilde{\alpha}_j^{species}$ to estimate the bacterial biomass $M_{bact} = M_{carb}/u$ from available carbohydrates. Note that with this calculation we assume that experimentally uncharacterized species present in a microbiota sample behave as the average of all characterized species. This is a first reasonable assumption to illustrate the sample-to-sample variation of fermentation product release as the relative abundance of the 22 characterized species commonly accounts for a high fraction of overall bacterial biomass (on average 57% for a collection of 219 microbiota samples from healthy individuals; see **Figure S2A** for coverage numbers). However, in subsets of samples, the total abundance of the experimentally characterized species is low, only accounting for a few percent of overall diversity (**Figure S2B**, right column). To better estimate the average rates from the experimental data we thus also performed the weighting calculation on coarser taxonomic levels with characterized species representing all species belonging to the same taxonomic group (TG). This approach is justified by the similarity in metabolism and fermentation pathways among closely related species (**Supplementary Text Section 1**), which we further confirmed in a bioinformatics study[25]. Specifically, we used the abundance data of detected species and their taxonomic classification to determine the fraction of biomass $\alpha_{TG}^{level}$ of all taxonomic groups which experimentally characterized strains represent for each of six specific taxonomic levels (phylum, class, order, family, genus, species). For example, on the genus level, eight taxonomic groups (genera) are represented by the 22 characterized strains and we calculated their abundance by accounting for all the detected species belonging to each taxonomic group (e.g. $\alpha_{Bacteroides}^{genus}$ is often the most abundant genus when analyzing microbiota samples from healthy donors, see **Figure 3A**). With this abundance, we then calculated the average per biomass excretion and uptake similarly as before. For example, for the genus level, we calculated the excretion as

$$\epsilon_i^{genera} = \sum_{TG \in represented\ genera} \epsilon_{i,TG} \cdot \tilde{\alpha}_{TG}^{genera}$$

With $\tilde{\alpha}_{TG}^{genera} = \alpha_{TG}^{genera} / \sum_j \alpha_{TG}^{genera}$ denoting the renormalized abundance. $\epsilon_{i,TG}$ denotes the typical excretion for the specific taxonomic group which we calculated by averaging (without weighting) over all the excretion rates $e_i$ from experimentally characterized strains belonging to this taxonomic group. Similarly, the uptake rate follows as

$$\mathcal{u}^{genera} = \sum_{TG \in represented\ genera} \mathcal{u}_{TG} \cdot \tilde{\alpha}_{TG}^{genera}$$

with $\mathcal{u}_{i,TG}$ denoting the typical uptake for the specific taxonomic group which we calculated by averaging (without weighting) over all the uptake rates $\mathcal{u}$ from experimentally characterized strains belonging to a specific taxonomic group.

The advantage of these calculations on a coarser taxonomic level (e.g. genus or family) is that the coverage of bacterial biomass becomes much higher. For example, for the genus level, on average a fraction of 81% percent of biomass is accounted for by this calculation when analyzing a collection of 219 microbiota samples from healthy individuals, with the fraction accounted for between 70% and 100% in almost all the samples (**Figure S2B**, middle panel). However, we also introduce the additional assumption that experimentally characterized strains represent all species of the same taxonomic group. This assumption, becomes imprecise on higher taxonomic levels, as species belonging to the same taxonomic group then come with a higher variety of metabolic strategies. As a standard, we thus do these calculations at the genus level as this level ensures a high coverage of biomass while still differentiating between metabolically diverse taxonomic groups of the phyla Bacteroidota and Bacillota. These phyla include the genera Phocaeicola and Bacteroides with the Prevotella and Bacteroides species known to differ in some aspects of their central metabolism (**Supplementary Text Section 1**). Another important example is the diversity of the family Lachnospiraceae, which includes butyrate producers but also acetogens.

To illustrate the variation of the fermentation product release with different diet conditions while keeping the microbiome constant (**Figures 2 and 4**), we chose a reference metagenomic dataset for the main text and specifically performed the rate calculations outlined above using average species abundance numbers observed across 219 samples from healthy adult donors. To analyze the variation of fermentation product release with microbiota composition while keeping diet conditions constant (**Figure 3**) we performed the calculations for each sample in the same reference metagenomics dataset of 219 samples. Resulting variations in fermentation product release for other collections of microbiota samples and with different dietary, age, and health parameters are presented in the **Data S1**. To further illustrate how the weighting calculations on different taxonomic level impacts the estimated release of fermentation products, we calculated the harvest for the reference microbiome collection also for taxonomic levels other than genus (**Figure S6**). All calculations to obtain the weighted per biomass uptake and excretion rates, as well as the resulting daily fermentation product release were performed with Python scripts which are available in the GitHub repository. Calculations to obtain the daily release of fermentation products based on diet parameters, given weighted uptake and excretion rates determined as described above, are introduced next.

### 3. Calculation of fermentation product release

In this section, we introduce detailed calculations to estimate the daily release of fermentation products depending on dietary and fecal weight numbers. The corresponding numbers for the British reference scenario are shown in following table.

**Major observables used in estimation for British Diet Reference Scenario.**
*variation based on observed variation reported in [26]
**variation based on reported variation in [27]

| Observable | Symbol | Value | Reference |
|---|---|---|---|
| Fecal wet weight | $M_{feces,wet}$ | $118g$ | [26,28] |
| Fraction dry mass | $\alpha_{dw}$ | $0.25$ | [26,28], see Figure S5 |
| Fecal dry weight* | $M_{feces,dry}$ | $29.5 \pm 6.8\ g$ | this study |
| Fraction of bacterial dry mass per fecal dry mass | $\alpha_{bact}$ | $0.55$ | [28] |
| Microbiota-available carbohydrates (standard estimation)** | $M_{Carb}$ | $35.7 \pm\ 9\ g$ | [27] |

#### 3.1. Estimation via feces

The starting point for this estimation is the daily amount of fecal wet weight secreted, $M_{feces,wet}$. With the fraction of fecal dry mass per wet mass, $\alpha_{dw}$, the daily amount of fecal dry weight excreted follows as

$$M_{feces,dry} = \alpha_{dw} \cdot M_{feces,wet}$$

Given the fraction of bacterial biomass per fecal dry mass, $\alpha_{bact}$, the daily amount of bacterial dry mass released then follows as

$$M_{feces,bact} = \alpha_{bact} \cdot M_{feces,dry}$$

The total daily amount of fermentation products released to support the growth of this biomass follows as

$$FP_{tot} = \sum_i FP_i = \sum_i \epsilon_i^{genera} M_{feces,bact}$$

with the sum over all major fermentation products the gut bacteria release, $FP_i$, with $i = \{acetate, butyrate, formate, lactate, propionate, succinate\}$. $\epsilon_i$ indicates the excretion of fermentation product $i$ per bacterial biomass). We obtain these numbers from the experimental characterization of 22 strains in pure cultures (**Figure 1**) weighted by the relative abundance of species in the microbiota (**Supplementary Text Section 2** above). We can further simplify this formula and define a total excretion coefficient $\epsilon_{tot} = \sum_i \epsilon_i$, such that the total amount of excreted fermentation products follows as

$$FP_{tot} = \epsilon_{tot} \cdot M_{feces,bact}$$

##### 3.2. Estimation via carbohydrates

The starting point for this estimation is the amount of carbohydrates which is available for bacterial digestion every day, the mass of microbiota available carbohydrates $M_{MACs}$. This number can come from a direct characterization of luminal content or by estimating the diet-dependent mapping between consumed nutrients and those reaching the large intestine (see **Supplementary Text Section 4** below), which specifies the fraction of total carbohydrate that passes the large intestine and the fraction which will be used by the microbiota. Assuming bacteria utilize all these available carbohydrates for fermentative growth, the bacterial biomass produced per day follows as:

$$M_{bact} = M_{MACs}/u$$

Here, $u$ indicates the per biomass uptake of carbohydrates, determined from measured uptake rates, again weighted by species abundance (**Supplementary Text Section 2**). Similar to the estimation via feces, the total daily amount of fermentation products released to support the growth of this biomass then follows as:

$$FP_{tot} = \sum_i FP_i = \sum_i \epsilon_i \cdot M_{bact}$$

with $i = \{acetate, butyrate, formate, lactate, propionate, succinate\}$ denoting different fermentation products and $\epsilon_i$ indicating the excretion of fermentation product $i$ per bacterial biomass, weighted by species abundance.

##### 3.3. Estimation via theoretical ATP yields

McNeil [29] previously built on ATP requirements to estimate the amount of fermentation products released by the human gut microbiota. Here, we repeat this estimation with updated numbers, starting with the bacterial mass in feces (16 g/day for the British reference diet; see **Figure 2A**). With well estimated ATP requirements for the biomass synthesis of fast-growing bacteria (35 mmol ATP per g dry biomass; **Supplementary Text Section 1.2**), 560 mmol ATP is needed every day to produce this biomass. With fermentation leading to the excretion of about 0.5-1 fermentation products per ATP synthesized (not including formate, **Supplementary Text Section 1.2**), the total fermentation products released would amount to 280-560 mol/day. The number is in line with our two estimates for the British reference diet via feces and via carbohydrates (**Figure 2AB**). Furthermore, with 2-4 ATP obtained per glucose molecule, 25.4 - 50.8 g of carbohydrates (such as fibers and starches) are needed to support growth, in agreement with the microbiota available carbohydrates estimated for the British reference diet (**Figure S4E-J**). Still, this ATP-based estimation comes with a substantial amount of uncertainty as (i) the amount of fermentation products released per ATP and (ii) the amount of ATP generated per glucose vary by about a factor 2, depending on the involved pathways. In addition, the 35mmol ATP/g energy requirement has been established well for *E. coli* [23], but it is less clear how this number changes across bacterial species and reported ATP requirements differ substantially between studies. To circumvent this uncertainty, we did not utilize ATP requirements for the estimations of the total fermentation product release we present in the main text. Rather, we build on our experimental characterizations of 22 highly abundant members of the human gut microbiota.

Notably, in his original estimation, McNeil used numbers for bacterial mass (15-20 g/day) and ATP yields per glucose (5 ATP per glucose molecule) comparable to those used above for the British reference diet. However, McNeil assumed a substantially higher ATP requirement of 100 mmol ATP per g bacterial dry weight. This led to the estimation that 59-65g/day of carbohydrates are needed to support bacterial growth and that 500-600mmol/day of fermentation products are released in the process. These numbers are substantially higher (i) compared to our ATP-based estimation above and (ii) compared to our via-feces or via-carbohydrate estimations based on experimentally measured carbohydrate uptake and fermentation product excretion values (about 460 mmol fermentation products/day). Furthermore, the estimated carbohydrate amount needed to fuel growth is not in line with the microbiota available carbohydrates estimated for the British diet (about 36g/day instead of 59-65g, **Figure S4E-J and Supplementary Text Section 4**). This discrepancy stems almost exclusively from the different assumptions on ATP requirements for biomass accumulation used by McNeil. Notably, McNeil based this number on data from ruminal bacteria characterized in continuous culture setups (chemostat) at very slow growth rates (<0.1 1/h) [30,31]. For slow growth under nutrient limiting conditions, maintenance energy can substantially contribute to the total ATP requirement, which leads to a larger ATP requirement for biomass synthesis than in conditions of fast growth. Notably, these slow growth conditions are likely not relevant for the human gut. Particularly, following nutrient influx from the small intestine [32], bacterial biomass in the large intestine is mostly accumulating in nutrient-replete conditions that allow for fast growth[33]. Notably, due to the higher ATP requirements, McNeil also estimated that the uptake of fermentation products by the human host covers 6-9% of the host's energy expenditure. Our different estimations revise this number, with a lower energy fraction of about 5% for diets resembling the British reference diet. Fractions up to 10% are only reached for non-Western diets extremely rich in complex carbohydrates (**Figure 6**).

#### 4. Mapping of nutrition data to microbiota-available carbohydrate

Nutritional data is commonly reported as a breakdown of total carbohydrate, protein, fat, and dietary fiber, as well as the amount of sugar consumed by a person per time period. We use data from the 1976 report on household food consumption and expenditure of the National Food Survey Committee in the United Kingdom [27] to estimate the microbiota-available carbohydrates for the British reference scenario as follows. First, we assume all consumed sugar is absorbed in the small intestine, and thus lost to the microbiota. For glucose, uptake rates as high as 550mmol/h have been measured in humans [34,35], a number which is sufficient to explain the uptake of all glucose even in meals containing large amounts. Lower numbers of around 100mmol/h are reported for the hydrolysis and uptake of sucrose, the main sugar component in human nutrition [36] but these numbers are still very high compared to typical sugar consumption (e.g. about 59g or 172.4mmol sucrose per day for British reference diet). The situation might change in cases where most sugar is taken up in more poorly absorbable forms such as fructose, where uptake rates are lower and transporters are easily saturated [37–39].

Second, we assume a constant fraction of the non-fiber, non-sugar carbohydrates to be available to the gut microbiota. This fraction of dietary carbohydrates mostly contains starches. Many starches can be enzymatically cleaved by host enzymes into glucose which is taken up in the small intestine and constitutes an important source of calories for the human body. Even though the molecular components of starches are well-defined, starch breakdown is not. Different types of starch molecules are degraded at different speeds, depending on the folding and packaging of the macromolecule, water content, molecular structure, and temperature in complex ways [40,41]. A fraction of the starches is completely inaccessible to host enzymes (the resistant starch fraction). Accordingly, the fraction of starches that pass through the small intestine into the large intestine has been reported to vary between 0% (pure soluble starch) and 96% (banana flour, [40]), with large variations, also depending on the type of food preparation [42–44]. However, when considering staple foods, such as boiled potato and white bread, which account for most starch in the British reference diet, the numbers reported for the fraction of starch reaching the large intestine vary only between 10% and 15% [40]. For the analysis presented in the main text we assumed a fraction of 13% of total starch to be available to the microbiota. To further illustrate the possible variation of fermentation product release with starch availability, we further discuss the obtained distribution values of fermentation products for a lower and higher passage of starches (**Figures S4G-J and S8)**.

Third, we assume that a constant fraction of dietary fiber is accessible to the gut microbiota. In contrast to starches, almost all dietary fibers reach the gut microbiota. However, not all of it can be broken down by bacterial enzymes. This fraction can depend on the composition of the microbiota, as some fiber can only be broken down by highly specialized bacteria [45,46]. For example, some common plant fibers such as cellulose and lignin are often inaccessible to most members of human gut microbiotas [47]. They contribute to fecal bulk as non-bacterial mass and have a role in water retention in the colon, but do not contribute to bacterial fermentation. For our analysis, we assume a fraction of 50% of dietary fiber to be accessible to bacterial enzymes and thus serve as a nutrient source for the gut microbiota. In **Figures S4** and **S8**, we analyze how our results would change with lower (25%) and higher (75%) digestion fractions.

Importantly, fiber consumption can further impact bacterial fermentation via its strong effect on transit times. Given the high water-binding capacity of fiber, high fiber consumption can substantially increase the bulk volume that passes the large intestine each day. Given a limited variability of the intestinal volume, fiber consumption thus leads to a strong increase in fecal wet weight and a strong decrease in transit time [48]. This is confirmed, for example, by a strong anticorrelation between transit time and fecal wet weight (**Figure S9D**). Importantly, for very high fiber consumption (and fecal wet weight), transit times can drop substantially below 24 hours. With 2-6 hours required for food to reach the large intestine, the biomass remains in the large intestine for only a very limited amount of time. Particularly for harder-to-digest carbohydrates, such short transit can severely reduce the efficiency with which bacteria digest carbohydrates. In our estimations, we do not explicitly model this dependence, but instead note that our analysis provides an upper boundary estimate, particularly for high fiber consumption.

### 5. Carbon sources not derived from dietary carbohydrates

In this study we mostly focus on dietary carbohydrates as the most abundant nutrient source promoting microbial growth along the large intestine. However, dietary protein reaching the large intestine and mucus released by epithelial cells can serve as alternative nutrient sources for the microbiota and also support fermentation. In this section, we analyze the quantities of these nutrient sources and the stoichiometry of protein fermentation. We show that the catabolic utilization of proteins and mucus contributes much less to the total amount of fermentation products produced by the gut microbiota than the catabolic growth on dietary carbohydrates.

#### 5.1. Protein-derived fermentation products

Proteins are an essential part of the human diet and constitute a major weight fraction of the food consumed (**Figures S4E, S7EG**). Most of the dietary protein is enzymatically digested in the stomach and small intestine to subsequently be taken up in the form of amino acids along the small intestine. Correspondingly, only a small fraction of consumed proteins reaches the large intestine. Controlled dietary studies that quantify the composition and turnover of intestinal fluids show this very well: for the protein load of the British reference diet (~70g/day), 6-7g of proteins, or around 10% of dietary protein, reach the large intestine each day, mainly in the form of peptides and not free amino acids[49]. Notably, a substantial fraction of the peptides detected might come from mucin and not the diet (see also below). However, to estimate an upper bound for the fermentation products derived from dietary protein, we here assume a protein load of about 10% of the dietary intake, $\frac{d}{dt} M_{prot,\ LI}^{BRD} = 7\ g/day$ for the British reference diet.

Since most gut bacteria reside within the large intestine and oxygen facilitates microbial respiration in the lumen of the small intestine, we consider only this protein load in our calculation of fermentation products. How many fermentation products are released by bacteria utilizing these peptides? This amount depends on two sequential processes. First, proteins need to be digested to gain free amino acids or very short peptides. Second, free amino acids need to be fermented.

<u>Step 1 - Protein digestion to amino acids:</u> Different gut bacteria differ strongly in their capability to digest proteins [50]. Overall protein digestion thus depends not only on the total amount of microbial biomass in the gut but also on the composition of the gut microbiota. Following observed digestion rates[50] and microbial biomass, we estimate that the time to digest ~10g proteins is substantial (~1 day), roughly comparable to the time biomass resides within the large intestine, which itself is well approximated by the transit time (0.5-3 days, **Figure S9D**). Thus, depending on transit times and microbiota composition, the microbiota should commonly not be able to efficiently digest all proteins reaching the large intestine. This is in line with observations showing that metabolites indicative of protein catabolism correlate positively with gut transit times[51]. More detailed studies are needed to understand the intricacies of protein digestion and its dependency on microbiota composition, transit time, and other factors. Here, we, therefore,

provide only an upper bound estimation of fermentation product release by assuming that proteins reaching the large intestine are fully digested by the microbiota, bearing in mind that this likely results in a substantial overestimation. For the British reference diet, we estimate the daily amount of amino acids available for gut bacteria to utilize to be

$$\frac{d}{dt}N_{AA}^{BRD} \approx \frac{\frac{d}{dt}M_{prot,LI}^{BRD}}{M_{AA}} \approx 60\frac{mmol}{day}$$

$M_{AA} = 110g/mol$ denotes the average molecular weight of amino acids.

<u>Step 2 - Amino acid utilization:</u> When amino acids are taken up by the bacteria, the release of fermentation products depends on the cellular use of amino acids. No fermentation products are released when amino acids are used in peptide translation and incorporated into newly synthesized proteins. In contrast, fermentation products are released when amino acids are used catabolically to provide energy to the cell. Since a large fraction of microbial mass is protein mass (commonly 40% or more of the cellular dry mass), and growing microbes thus need to synthesize large amounts of proteins, a substantial fraction of the amino acids should end up in cellular proteins and thereby not contribute to the excretion of fermentation products. However, as an upper bound estimation, we assume here that all amino acids reaching the colon are used catabolically. The exact type and amount of fermentation products released depends on the specific amino acids and metabolic reactions [52]. However, Stickland fermentation, the coupled use of two amino acids, with one serving as electron donor and the other as electron acceptor, seems to be the most common way to catabolize amino acids: Stickland fermentation is typically faster than uncoupled amino acid fermentation and the preferred process when amino acids are available as electron acceptors [52,53]. In addition, at least for in-vitro experiments with rumen communities, chemical inhibition of Stickland fermentation inhibits amino acid deamination, the first step of amino acid fermentation [54,55]. A Stickland oxidation-reduction pair generally releases 0.5 ATP and 1 fermentation product. Notable, depending on the specific pairs of amino acids involved, the types of fermentation products released vary. Many are branched chain fatty acids and not the fermentation products released when carbohydrates are utilized, with potential different effects on the host[51]. In our calculations, we do not distinguish between these types of fermentation products. For the British reference diet, the upper bound for protein-derived amino acid fermentation we obtain is therefore:

$$\frac{d}{dt}N_{FP}^{BRD} = 1 \cdot \frac{d}{dt}N_{AA}^{BRD} \approx 60\frac{mmol}{day}$$

#### 5.2. Mucin-derived fermentation products

We similarly determine the contribution of mucin digestion to microbiota-derived fermentation products. We first estimate the daily supply of mucin by the host, and then its subsequent metabolic utilization by bacteria.

<u>Step 1 - Daily mucin supply:</u> Mucin is released in substantial quantities from the mucus layers of the small and large intestines. The amount released in the small intestine can be estimated from

the characterization of lumen content enabled via tubing-assisted sampling. Assuming all hexosamines measured come from mucin, the upper bound of mucin released by the small intestine and entering the large intestine is about $\frac{d}{dt} M_{mucin,SI}^{overest} = 3 - 4 \frac{g}{day}$ [56]. Direct measurements of mucin released in the large intestine are not available. However, we can estimate the release using well-characterized numbers on mucus thickness, content, and turnover: the mucus coating the epithelium of the large intestine in mice and rats consists of two layers, a dense inner layer with a thickness of $50 - 100\mu m$ and a looser outer layer with a thickness of $100$ to $700\mu m$ [57]. We assume $w_{muc,LI}^{dense} = 100\mu m$ and $w_{muc,LI}^{loose} = 700\mu m$ as the thickness of the dense and loose mucus layers, respectively, and further consider the typical length ($l_{LI} = 150cm$) and diameter ($d_{LI} = 4cm$) of the adult human large intestine to determine the total mucus volume in the large intestine for both the dense and loose layers, $V_{muc,LI}^{dense} = \pi \cdot d_{LI} \cdot l_{LI} \cdot w_{muc,LI}^{dense} \approx 19ml$, and $V_{muc,LI}^{loose} = \pi \cdot d_{LI} \cdot l_{LI} \cdot w_{muc,LI}^{loose} \approx 132ml$. The mucus content in mucus layers has been reported to range between $\alpha_{mucin} = 0.2 - 5\%$[58], and the volume of gastrointestinal mucus has been reported to expand by a factor 3-4 at the transition between dense and loose layers [59]. We thus take the percentage of mucin in the dense layer to be $\alpha_{mucin}^{dense} = 5\%$ and the percentage of mucin in the loose layer to be $\alpha_{mucin}^{loose} = \frac{5}{3}\%$. With these numbers we calculate the amount of mucin present in the mucus layers of the large intestine as: $M_{mucin,LI} \approx V_{muc,LI}^{dense} \cdot \rho_{water} \cdot \alpha_{mucin}^{dense} + V_{muc,LI}^{loose} \cdot \rho_{water} \cdot \alpha_{mucin}^{loose} \approx 3.1\ g$.

The daily release of mucin follows from the mucus amount and the turnover of the mucus layer with new mucin continuously replacing old mucin, $\frac{d}{dt} M_{mucin,LI} = \frac{M_{mucin,LI}}{2 \cdot t_{half}}$. Here, $t_{half}$ denotes the half-life time characterizing this turnover. This number has been quantified well in mice using isotope labeling techniques with $t_{half} \approx 4\ days$ for the colon of conventional mice, notably longer than the half-life time within the small intestine[60]. With this number, the daily mucin release in the large intestine is $\frac{d}{dt} M_{mucin,LI} \approx 0.4\ g/day$, leading to a total release of $\frac{d}{dt} M_{mucin} = \frac{d}{dt} M_{mucin,LI} + \frac{d}{dt} M_{mucin,SI} \approx 4.4\ g/day$.

<u>Step 2 – Mucin utilization by bacteria:</u> Mucins are glycoproteins that contain carbohydrates and peptides with a well-known mass fraction for the most abundant mucin MUC2 (20% proteins and 80% carbohydrates)[57]. The utilization by bacteria relies on degradation capabilities, which can strongly vary between strains and thus with microbiota composition. To obtain an upper estimation, we assume here that all mucin is completely digested, leading to a total amount of mucin-derived peptides and carbohydrates:

$$\frac{d}{dt} M_{mucin,carbohydrates} \approx 4\ g/day\ ,$$

$$\frac{d}{dt} M_{mucin,c\ proteins} \approx 1\ g/day.$$

From these numbers, we estimated an upper bound on the mucin-related release of fermentation products, assuming all mucin-derived peptides and carbohydrates are utilized for fermentation and following the estimations shown before for carbohydrates (**Supplementary Text Section 3.2**) and proteins (**Supplementary Text Section 5.1**).

$$\frac{d}{dt} N_{mucin,FP} \approx 2\, mmol\ glucose\ eq./day\ ,$$
$$\frac{d}{dt} M_{proteins,FP} \approx 9\, mmol\ AA/day.$$

##### 5.3. Contribution of protein and mucin fermentation to overall release of fermentation products

Compared to dietary carbohydrate-derived fermentation products, the release of protein and mucin-derived fermentation products are low. For the British reference diet, the comparison is shown in **Figure 2D,** with the total amount of fermentation products being maximally 25% higher when taking the upper bound estimates of protein- and mucin-derived fermentation products. Furthermore, because of our consistent overestimation of the role of mucin and dietary protein digestion, we are likely overestimating this difference very strongly. When taking dietary proteins into account, the estimated upper bound difference in the fermentation product release within the US population (**Figure S8D**) and for the Hadza (**Figure S8H**) is similar and also small. We conclude that utilization of dietary protein and mucus as substrates for fermentation increase the release of fermentation products, but in comparison to complex carbohydrate fermentation, this increase is low, even for our clear overestimations of mucin and protein digestion.

### 6. Most released fermentation products are taken up by the host

To estimate the amount of fermentation products that the host obtains from growing bacteria, we need to estimate the fraction of fermentation products that are taken up by the host. As we outline in the main text, we estimate this fraction to be very high, and we therefore take the produced amount of fermentation products as a good (upper bound) estimate for the consumed fermentation products by the host. In the following, we discuss the underlying reasons. Besides consumption by the host, fermentation products might specifically be lost via feces or consumed by cross-feeding microbes. In addition, the turnover of fermentation products might also be shaped by the recycling of bacterial biomass. In the following we discuss these three different possibilities separately.

Loss of fermentation products via feces is negligible: Concentrations of fermentation products in feces have been measured, and the summed concentration of all major fermentation products rarely exceeds about 100mM [61]. With an average fecal wet mass of 118 g/day for a Western diet [26], this corresponds to approximately 10 mmol of fermentation products which are secreted via feces each day. With the human gut microbiota releasing approx. 500 mmol of fermentation products each day (British reference scenario, **Figure 2**), about 2% or less of the fermentation products leave the body via feces. Accordingly, most fermentation products remain in the host. These numbers also emphasize that measurements of fermentation products in feces are a bad proxy for fermentation products released by the microbiota.

Cross-feeding likely does affect the total amount of fermentation products released by the gut microbiota in major ways: Various forms of cross-feeding within the gut microbiota, with members of the microbiota consuming metabolites released by other members, has been demonstrated[62–66]. The often discussed methanogens and sulfate reducers utilize H2 and CO2 or sulfate, respectively, without directly affecting the concentrations of fermentation products in the environment[67,68]. There are, however, cross-feeding processes that involve the uptake of major fermentation products, including lactate and acetate [6,62]. Importantly, however, as these processes also lead to the release of other fermentation products, such as butyrate, this type of cross-feeding has only a limited effect on the net release of fermentation products. Furthermore, the primary example of this type of cross-feeding is the incorporation of acetate into butyrate, reported for some of the most abundant gut microbiota members, including *A. rectalis* and *R. intestinalis* [6]. Our measurements account for this process as YCA medium contains acetate and our experimental data thus reflect net changes in this metabolite. Lactate cross-feeding in *A. caccae* and *A. hallii* is another important example of this type of cross-feeding. It has been shown to depend on pH and carbohydrate availability [69]. Determining the extent to which this cross-feeding shapes the fermentation product balance in the gut will be important future work. Another type of cross-feeding is acetogenesis. Acetogens can produce additional acetate from CO2 and H2 independent of carbohydrate or short chain fatty acid supply. In our data, we see this reflected in the carbon balance of the well-described acetogen *B. hydrogenotrophica* (**Figure 1F**). However, this process likely has only a minor effect on the fermentation product balance because of the very low abundance of *B. hydrogenotrophica* (<0.1%). In conclusion, cross-feeding likely affects the relative abundances of fermentation products (with potential consequences for the local environment and uptake dynamics), but its effects on the total fermentation product harvest is limited.

Recycling of bacterial biomass is unlikely to substantially impact the harvest of fermentation products: Bacterial biomass contains sugars, proteins, and other potential nutrient sources [70]. For these to become available for other bacteria, bacterial cell components would have to leak into the gut lumen. This probably happens to some extent, but this effect is likely small and negligible when calculating the net turnover of biomass and fermentation products. This is in part because the death of bacteria is a relatively slow process, happening for example at a rate of about 0.4 $d^{-1}$ for starving *E. coli* cultures in aerobic batch cultures [71], comparable or longer than the transit time biomass remain in the large intestine. But even if cells lyse, the effect on fermentation is likely limited. For example, the addition of supernatant from dying cultures can halt the death process in batch cultures, but it does not support biomass growth [71].

Taken together, these considerations suggest that most released fermentation produced are taken up by the human host. Furthermore, our analysis itself confirms a minor role for biomass recycling and cross-feeding as the total bacterial mass lost each day in feces is very close to the biomass that can be generated based on the amount of carbohydrates available to the microbiota.

### 7. Mouse estimations

#### 7.1. Estimation via feces

To estimate the contribution of the microbiota to energy homeostasis in mice we first followed a similar calculation as for humans using the via-feces estimation. We start with data for conventionally colonized mice fed ad libitum [72]. The daily fecal dry mass these animals release is

$$M_{fec}^{mouse} = 0.82 \pm 0.09\ g/day$$

The variation here describes the SD of the reported variation in fecal weight. The microbial density in mouse feces lies within the range of $10^{11}$ and $10^{12}$ cells per gram [73,74]. As we are not aware of more accurate measurements, we assume a similar fraction of the fecal dry matter to be bacteria as in humans (**Figure S4A-D**), about 50%, leading to an excreted bacterial mass of

$$M_{feces,bac}^{mouse} \approx 0.41 g/day$$

As described above (**Supplementary Text Section 3**), we calculate the amount of fermentation products as

$$FP_{tot}^{mouse} = \sum_i FP_i^{mouse} = \sum_i \epsilon_i M_{feces,bact}^{mouse} \approx 11.8 \pm 1.3\ mmol/day$$

As for humans, this translates into an energy contribution as follows (see **Supplementary Text Section 8** below)

$$E_{ferm}^{mouse} = \sum_i H_i \cdot FP_i \approx 10.6 \pm 1.2\ kJ/da$$

While variations remain, indirect calorimetry measurements for these mice indicate a daily energy expenditure of $E_{exp}^{mouse} \approx 38{,}0\ kJ/day$ [72]. With this, the bacterial contribution to energy homeostasis in mice is approximately,

$$\alpha_{energy}^{mouse} = \frac{E_{ferm}^{mouse}}{E_{exp}^{mouse}} \approx 28 \pm 3\%$$

To validate these results, we further use an alternative way to arrive at this number which is based on the comparison of germ-free and conventionally colonized mice.

##### 7.2. ESTIMATING MICROBIAL ENERGY OUTPUT BY COMPARING ENERGY RETENTION IN GERM-FREE AND COLONIZED MICE

As a second way of estimating the bacterial energy contribution in mice, we compare energy retention from food in conventionally colonized (CC) and germ-free (GF) mice, feeding ad libitum on chow. It is well known that food consumption is substantially higher in GF animals [72,75,76], which is consistent with a substantial bacterial contribution to energy homeostasis.

To estimate the energy fraction obtained from the microbiota, we start with the energy of daily consumed food determined in germ-free and conventional mice, as determined via calorimetry measurements [72]:

$$H_{food}^{CC} = 54.5 \pm 8.5 \frac{kJ}{day}$$

$$H_{food}^{GF} = 62.7 \pm 8.9 \frac{kJ}{day}$$

Here and in the following, the provided ranges denote variations (SD) based on observed variations in food intake. In line with the hypothesis that GF mice cannot utilize dietary polysaccharides, the energy content of feces also varies.

$$H_{feces}^{CC} = 13.6 \pm 1.8\ kJ/day$$

$$H_{feces}^{GF} = 25.0 \pm 4.6\ kJ/day$$

Notably, the total energy mice extract from food each day (i.e., the difference between energy content in food and feces) is highly comparable between conventional and GF mice,

$$E_{extr}^{conv} = H_{food}^{CC} - H_{feces}^{CC} \approx 37.6\ kJ/day$$

$$E_{extr}^{GF} = H_{food}^{GF} - H_{feces}^{GF} \approx 40.9\ kJ/day$$

indicating that mice fed ad libitum adjust their food intake to match the energy they need independent of the source of this energy. With these numbers, we can calculate the efficiency of energy extraction from both colonization states:

$$\frac{E_{extr}^{CC}}{H_{food}^{CC}} \approx 75 \pm 12\%$$

$$\frac{E_{extr}^{GF}}{H_{food}^{GF}} \approx 60 \pm 9\%$$

This indicates a microbiota-derived energy gain in conventionally colonized mice of

$$\Delta E_{microbiota}^{mouse} = \left(\frac{E_{extr}^{CC}}{H_{food}^{CC}} - \frac{E_{extr}^{GF}}{H_{food}^{GF}}\right) \cdot H_{food}^{CC} \approx 8.1 \pm 1.8\ kJ/day$$

Accordingly, a fraction of

$$\alpha_{energy}^{mouse} = \Delta E_{microbiota}^{mouse} / E_{extr}^{CC} \approx 21 \pm 5\%$$

of the daily energy expenditure $E_{extr}$ is supplied by the microbiota. This number is comparable to the estimation obtained via fecal microbial counts and the energy of fermentation products presented before ($28 \pm 3\%$).

#### 7.3. Estimating relevant amounts of fermentation products in humans and mice

According to our estimates, colonized mice typically gain substantially more of their daily energy need from fermentation products derived from their gut microbiota than humans (**Figure 6**). In absolute numbers, their fermentation product harvest amounts to around 12mmol/day (fecal estimate, see **Supplementary Text Section 8**), which is roughly a factor 40 lower than what we calculated for humans (470mmol/day, fecal estimate, see **Figure 2A(iii)**). However, assuming a typical body weight of 30g for a mouse and 70kg for a human, the body weights of the two organisms differ by a factor of roughly 2300. Normalizing by weight thus yields a systemic fermentation product harvest of approximately 400mmol/kg/day for mice and 7mmol/kg/day for humans.

In addition to systemic effects such as behavioral modulation, fermentation products have been shown to have effects locally at the mucosal surface of the intestinal tract [77–81]. The local fermentation product harvest is different from the systemic one, because the relative large intestinal surface area is much greater in mice than in humans (0.03m$^2$/kg for a 70kg human [82], 2.3m$^2$/kg for a 30g mouse [83]). Taking into account this difference in surface areas, the per-area values we obtain for the fermentation product harvest in mice and humans are much closer to each other (174mmol/m$^2$/day for mice, and 224mmol/m$^2$/day for humans). This difference in relative large intestinal surface area suggests an evolutionary adaptation of uptake surface to the larger fermentation product dependence in mice. Assuming that the dose dependence of mucosal immune modulation is comparable between mice and humans, the difference in the effect of fermentation products between the two organisms is likely small in this respect.

### 8. Estimation of energy contribution

We estimate the energy content of the fermentation products released using their combustion enthalpies [84]. The energy content of all fermentation products (except formate, for which we are not aware of any evidence that it is used for energy generation in the host) released each day follows from the release of each fermentation product and the corresponding enthalpy $H_i$ listed in **Table S1**.

$$E_{ferm} = \sum_i H_i \cdot FP_i$$

Most of these fermentation products are taken up by the host (**Supplementary Text Section 6**) where they are subsequently used in different parts of the body for respiration [85,86]. To assess the relative contribution of this process to the total daily energy expenditure $E_{exp}$ we calculate the ratio,

$$\alpha_{energy} = \frac{E_{ferm}}{E_{exp}}$$

For the British reference diet we used reported numbers, $E_{exp} \approx 2275\ kcal\ (9.52\ MJ)$ [87,88]. To analyze the variation within the US population, we use the estimations from the NHANES study reported for each study participant (**Figure S7D** [89]). For the global variation analysis via fecal weight, we assumed values reported for the British reference diet, $E_{exp} \approx 2275\ kcal\ (9.52\ MJ)$. For the Hadza, we calculated values based on the consumed carbohydrates, fats, and proteins reported for each month, on average $E_{exp} \approx 2240\ kcal\ (9.37\ MJ)$.
